# Liver microbiome composition associates with histological severity and PNPLA3 genotype in metabolic dysfunction-associated steatotic liver disease

**DOI:** 10.64898/2026.06.30.735597

**Authors:** Maria F Mascardi, Rosario Taussig, Ivan Privitera Signoretta, Barbara Suarez, Sebastian Marciano, Paola Casciato, Adrian Narvaez, Leila Haddad, Adrian Gadano, Alberto Penas-Steinhardt, Juan P Bustamante, Julieta Trinks

**Affiliations:** Instituto de Medicina Traslacional e Ingenieria Biomedica (IMTIB) - CONICET - Universidad Hospital Italiano - Hospital Italiano de Buenos Aires, Ciudad Autonoma de Buenos Aires C1199, Argentina; Consejo Nacional de Investigaciones Cientificas y Tecnicas (CONICET), Ciudad Autonoma de Buenos Aires C1425, Argentina; Facultad de Ingenieria, Universidad Austral, LIDTUA-CIC, Pilar B1630FHB, Buenos Aires, Argentina; Planta Piloto de Ingenieria Quimica (PLAPIQUI), CONICET, Universidad Nacional del Sur, Bahia Blanca B8000, Buenos Aires, Argentina; Seccion de Hepatologia, Servicio de Clinica Medica, Hospital Italiano de Buenos Aires, Ciudad Autonoma de Buenos Aires C1199, Argentina; Departamento de Investigacion, Hospital Italiano de Buenos Aires, Ciudad Autonoma de Buenos Aires C1199, Argentina; Departamento de Ciencias Basicas, Laboratorio de Genomica Computacional (GEC-UNLu), Universidad Nacional de Lujan, Lujan B6700, Buenos Aires, Argentina; Facultad de Ingenieria, Universidad Nacional de Entre Rios, Oro Verde CP3260, Entre Rios, Argentina

**Author notes:** **Corresponding author: Maria F Mascardi, MD, PhD(c),** Instituto de Medicina Traslacional e Ingenieria Biomedica (IMTIB)—CONICET—Universidad del Hospital Italiano (UHI)—Hospital Italiano de Buenos Aires (HIBA), Potosi 4240, Ciudad Autonoma de Buenos Aires (C1199ACL), Argentina.

**Keywords:** Metabolic dysfunction-associated steatotic liver disease, Patatin-like phospholipase domain-containing 3, Hepatic microbiome, Gut-liver axis, Shotgun metagenomics, Liver biopsy, Somatic mutational burden, Correlation networks

## Abstract

**BACKGROUND:** Metabolic dysfunction-associated steatotic liver disease (MASLD) is a systemic immunometabolic disorder rapidly increasing worldwide, affecting nearly 38% of adults. Gut dysbiosis and host genetic factors, such as PNPLA3 I148M variant, modulate disease development and progression. Through the gut–liver axis, increased intestinal permeability enables microbial translocation to the liver, promoting inflammation and metabolic disruption. However, the composition and functional potential of the hepatic microbiome remain poorly characterized. Understanding its relationship with histological injury and genetic susceptibility may provide novel mechanistic insights. We hypothesized that the hepatic microbiome composition and function are associated with histological severity and PNPLA3 genotype in this disease.

**AIM:** To characterize the hepatic microbiome and assess its association with histological severity and PNPLA3 genotype.

**METHODS:** This cross-sectional observational study included 30 patients with MASLD from a tertiary care hospital. Liver tissue underwent shotgun metagenomic sequencing. Histological severity was assessed using the NAFLD Activity Score (NAS). PNPLA3 genotype was determined by PCR. Differential abundance and functional enrichment analyses were performed using MaAsLin2. Somatic variants were identified using Mutect2. Correlation networks were constructed using Spearman’s correlation coefficients.

**RESULTS:** Patients with advanced histological injury (NAS ≥5) and PNPLA3 I148M carriers showed a trend toward higher somatic mutational load and a markedly reduced microbial abundance. Analyses revealed broad compositional shifts across bacterial, fungal, viral, and eukaryotic taxa, affecting both commensal and context-dependent pathobiont lineages. *Pseudomonas* species were enriched, whereas Siphoviridae phages were depleted in advanced disease and PNPLA3 I148M carriers. Functional analysis revealed enrichment of pathways related to nutrient transport and metabolic stress adaptation, while TonB-associated functions were enriched in advanced liver injury but depleted in PNPLA3 I148M carriers. Network analysis identified *Sphingomonas leidyi* as a keystone node associated with hexosamine metabolism. *Salmonella enterica* abundance positively correlated with somatic variant burden, suggesting a link between microbial signatures and genomic instability. Histological progression and the risk PNPLA3 genotype were accompanied by marked topological simplification, reflecting less resilient community structures.

**CONCLUSIONS:** The hepatic microbiome in MASLD is a low-biomass, polymicrobial ecosystem shaped by the host genetic background. Its functional activity, taxonomic composition and system architecture bidirectionally relate to liver DNA instability and the severity of histological damage.

**Core tip:** This study characterizes the multi-kingdom hepatic microbiome in MASLD using FFPE-derived metagenomics. We demonstrate that microbial abundance-including bacteria, fungi, protozoa, and viruses- significantly decreases with increased histological severity and the PNPLA3 risk genotype. Rather than global diversity shifts, results showed that disease progression could be linked to specific functional adaptations and simplified microbial network connectivity. In addition, we described associations between specific taxa and somatic mutational burden, suggesting a link between microbial signals and genomic instability. These findings indicate that changes in the liver microbiome as a whole, rather than specific taxonomic modifications, influence MASLD pathophysiology.

## INTRODUCTION

Metabolic dysfunction-associated steatotic liver disease (MASLD) represents the hepatic manifestation of a systemic immunometabolic disorder, clinically defined by the presence of hepatic steatosis alongside at least one cardiometabolic risk factor ^[1]^. This updated nomenclature more accurately reflects the condition’s intrinsic link to systemic metabolic dysregulation, distinguishing it from its predecessor, non-alcoholic fatty liver disease (NAFLD) ^[2]^. The global health burden of MASLD is substantial and growing: its prevalence has surged from an estimated 14-27% a decade ago to 38% of the current adult population, mirroring the rising rates of obesity and type 2 diabetes ^[3]^. The clinical significance of MASLD lies in its potential progression to metabolic dysfunction-associated steatohepatitis (MASH), an aggressive inflammatory state that drives fibrosis, cirrhosis, and hepatocellular carcinoma, positioning it as a leading cause for liver transplantation ^[4]^.

The pathogenesis of MASLD is multifactorial, involving a complex interplay of genetic predispositions and environmental factors ^[5]^. While genome-wide association studies have linked several host gene variants to the disease, the patatin-like phospholipase domain-containing 3 (*PNPLA3*) I148M variant (rs738409) stands as one of the most potent genetic risk factors for MASH and advanced fibrosis ^[6]^. However, recent studies recognize MASLD as an eco-biological disease, where liver disease risk is not only shaped by host genetics and environment, but also by the ecological configuration and functional outputs of the gut microbiome. This perspective redefines disease susceptibility as, in part, context-dependent and microbiota-mediated ^[7]^.

From this perspective, a central element in the pathogenic process of MASLD is the gut-liver axis, which describes the bidirectional physiological and anatomical communication between the gut, its microbiome, and the liver ^[8]^. In MASLD, the integrity of the intestinal barrier is often compromised, leading to increased permeability. This allows for the translocation of microbial-associated molecular patterns (MAMPs) from the gut lumen into the portal circulation. Upon reaching the liver, these molecules initiate and perpetuate a cycle of chronic inflammation and fibrogenesis, disrupt lipid and glucose homeostasis in the liver, or induce direct hepatotoxicity ^[9]^.This relationship is bidirectional, as the liver actively shapes the gut microbiome through the secretion of bile, IgA, and antimicrobial peptides into the intestine. Conversely, the diseased liver environment, characterized by altered bile flow and tissue hypoxia, can in turn reshape gut microbial communities, often favoring pathogenic configurations and creating a self-sustaining loop that drives disease progression ^[9,10]^. Notably, carriers of the *PNPLA3* risk allele GG have been shown to harbor a distinct liver microbiota signature, reflecting the complexity of MASLD and suggesting an intricate interplay between host genetics, the local microbiome, and metabolic inflammation as disease drivers^[11]^.

Despite this growing evidence, the composition and functional potential of the liver tissue microbiome itself, especially regarding its non-bacterial component, remain poorly characterized. Most studies that addressed the impact of the human microbiome on the pathophysiology of MASLD have focused exclusively on the gut microbiome or have relied on 16S rRNA gene sequencing of the liver microbiome, which has taxonomic limitations and only examined the bacterial component of the microbiome ^[11,12]^. Therefore, we aimed to comprehensively characterize the hepatic metagenome of MASLD patients. In addition, we evaluated its association with the severity of histological damage and with the *PNPLA3* risk genotype, providing novel insights into the role of the intrahepatic microbial ecosystem in the progression of MASLD.

## MATERIALS AND METHODS

This is an analytical observational study with cross-sectional design. It was performed in line with the principles of the Declaration of Helsinki. Approval was granted by the Ethics Committee of “Hospital Italiano de Buenos Aires”.

### Participant selection and sample collection

Thirty adult MASLD patients, who had been recruited in previous MASLD studies by our group ^[13]^, were retrospectively selected for this study based on the grade of liver injury. Half of the 30 recruited subjects were identified with a low degree of histological liver injury, forming the low NAS group (NAS score≤4), and the remaining 15 patients had a high degree of injury, forming the high NAS group (NAS score≥5). All of them were recruited at the Hepatology Unit of the “Hospital Italiano de Buenos Aires”. MASLD diagnosis was confirmed by liver biopsy, which along with standard medical practices allowed to rule out other liver diseases ^[14]^. Exclusion criteria for MASLD patients were: schistosomiasis, any liver disease other than MASLD, anticipated need for liver transplantation within a year or complications of end-stage liver disease, concurrent medical illnesses, and contraindications for liver biopsy. In addition, hepatitis B virus and hepatitis C infections were excluded in all MASLD patients by HBsAg, anti-HBc IgG and anti-HCV IgG analysis using the Architect Abbott system (Abbott Diagnostics, Wiesbaden, Germany).

Liver biopsy was obtained through the percutaneous approach under ultrasound guidance. Adequacy of liver biopsy samples was assessed by the length and diameter of its core. Therefore, a 15 to 25 mm length sample obtained by a 16-gauge needle was considered to be adequate. The severity of histological injury of the samples was assessed by an expert liver pathologist from the Pathology Unit of the hospital, based on components of the NAFLD Activity Score (NAS score). Tissue samples were fixed using formalin and subsequent embedding in paraffin wax (FFPE tissues) and preserved at room temperature.

In addition, each participant provided a fasting blood sample and underwent anthropometric assessments. Demographic information and medical history were also recorded. Blood samples were collected in tripotassium-EDTA tubes, promptly processed for plasma separation and isolation of peripheral blood mononuclear cells (PBMCs), and stored at −80°C.

### DNA extraction and determination of the PNPLA3 SNP rs738409

Genomic DNA was extracted from isolated PBMCs using the QIAamp® DNA Mini Kit (QIAGEN, GmbH, Hilden, Germany) following the manufacturer’s protocol. The PNPLA3-SNP was partially amplified via PCR using the primer pair 5′-CGA TCT AGC CCC TTT CAG TC-3′ (forward) and 5′-GCA GAT TAA GTG AAC CAG CC-3′ (reverse). The PCR protocol included 40 cycles, each consisting of denaturation at 94°C for 30 seconds, annealing at 62°C for 30 seconds, and extension at 72°C for 1 minute. The resulting PCR products, 668 base pairs in length, were visualized on 2% agarose gels using TBE buffer under UV light after electrophoresis at 100 V/min and staining with ethidium bromide.

Bi-directional sequencing of the PCR fragments was performed using the Big-Dye Terminator chemistry system (Applied Biosystems, Life Technologies Corp., Foster City, CA, USA), and sequence chromatograms were analyzed using BioEdit Sequence Alignment Editor (version 7.1.3.0).

### DNA extraction from FFPE tissues, shotgun metagenomic sequencing and DNA-seq data processing

Ten slices of 10 μm were subsequently sectioned from each FFPE tissue block and subjected to DNA extraction using QIAamp DNA FFPE Advanced UNG Kit (QIAGEN®). For shotgun metagenomic sequencing, libraries were prepared following the Illumina DNA Prep protocol (Illumina®). All procedures were performed under sterile conditions to avoid contamination. Once the libraries were completed, they were individually quantified using Qubit® 4.0 and Qubit^®^ dsDNA high-sensitivity (Thermo Fisher Scientific, Waltham, MA, EE.UU.). All libraries were then pooled into equimolar proportions and assessed on a Bioanalyzer 2100 with the High Sensitivity DNA kit (Agilent Technologies, Santa Clara, CA, EE.UU.). DNA sequencing was performed on the Illumina® NextSeq 550 sequencer in 2x150bp paired-end mode.

A total of 189,696,188 sequencing reads were generated. The raw reads were processed using Trimmomatic v.0.39 to remove adapter sequences and trim low-quality bases. After quality control, 162,838,445 reads (85.84% of the raw data) were retained. These reads were then aligned to an expanded human reference genome (db GRCh38) using Bowtie2 to filter out host-derived sequences. Ultimately, 1,206,693 reads (0.74% of the trimmed reads) passed the filtering step and were used for subsequent analyses.

### Taxonomic and functional characterization of liver metagenome

Short read alignment against the NCBI RefSeq non-redundant proteins database (v.feb2022) was performed using DIAMOND ^[15]^, followed by taxonomic and functional binning using MEGAN6-C.E (LCA parameters: minimal bit score 87, top percent 5, minimum number of assigned reads 3) ^[16]^. To reduce spurious assignments, reads classified under *Deuterostomia*, *Panarthropoda*, and *Viridiplantae* were excluded from downstream analysis. This filtering step is justified by the behavior of DIAMOND–MEGAN workflows, in which individual short reads may generate multiple significant alignments across distant taxa. MEGAN incorporates all such hits during LCA computation and propagates each read upward through all ancestral nodes, allowing low-frequency artefactual matches to inflate biologically implausible eukaryotic clades. Given the low microbial biomass of FFPE liver biopsies and the susceptibility of such samples to host carryover and environmental contamination, the higher eukaryotic taxa mentioned before typically reflect misclassification or exogenous DNA rather than authentic liver microbiome members. This approach follows MEGAN’s recommended contamination-filtering practices, which permit exclusion of entire taxonomic lineages in human-associated microbiome datasets ^[17]^.

Taxonomic composition was assessed at family, genus, and species level. For functional characterization, the predicted protein domains were mapped to the InterPro protein family database ^[18]^, whereas eggNOG database ^[19]^ mapping allowed for functional inference based on orthology.

All the obtained liver microbiome data was independently compared between groups regarding NAS score (NAS≥5 *vs*. NAS≤4) and PNPLA3 genotype (GG genotype *vs*. CC/CG genotype). Differential abundance and functional enrichment analyses were performed using the MaAsLin2 software ^[20]^. To account for differences in sequencing depth across samples, the "cumulative sum scaling" (CSS) normalization method was applied. For statistical comparisons between groups, the "negative binomial" model was employed, which is particularly suited for metagenomic count data as it accounts for overdispersion commonly observed in microbial datasets ^[20]^. Differences were considered significant if the variable (taxonomic or functional feature) was observed in at least 10 samples with a significance threshold at *q*<0.01.

Alpha diversity of species was calculated using Shannon and Chao1 indices and compared between groups using the Wilcoxon test. For Chao1 indices calculation, normalization on read counts was performed by means of rarefaction to the minimum sample sequencing depth using random subsampling without replacement. Beta diversity was evaluated based on Bray-Curtis distances and compared statistically using PERMANOVA. All diversity analyses were performed in R using the *vegan* package ^[21]^. In both cases, the significance threshold was set at *p*<0.05.

### Analysis of somatic mutational burden in FFPE liver biopsies

Due to the distinct aims of the metagenomic and somatic variant analyses, we applied different quality control pipelines optimized for each workflow. While the metagenomic pipeline aimed to remove human reads and retain microbial DNA, the somatic variant analysis required high-fidelity alignment to the human reference genome. Thus, raw reads intended for somatic variant calling were processed with KneadData v.0.6.1, included in the bioBakery3 platform ^[22]^, which integrates Trimmomatic v0.39 ^[23]^ for trimming adapters and low-quality bases, and Bowtie2 ^[24]^ for filtering out contaminant microbial sequences using the SILVA database ^[25]^.

High-quality reads were subsequently aligned to the human reference genome (GRCh38, obtained from https://storage.googleapis.com/gcp-public-data--broad-references/hg38/v0/Homo _sapiens_assembly38.fasta) using BWA-MEM ^[26]^. The resulting SAM files were converted to BAM format using Samtools ^[27]^, and downstream processing was performed with GATK v4.2 ^[28]^. This included the addition of read group information using AddOrReplaceReadGroups, sorting with SortSam, and BAM indexing with BuildBamIndex.

Somatic variants were called in tumor-only mode using Mutect2 ^[29]^, employing the af-only-gnomad.hg38.vcf.gz database as a germline resource (downloaded from https://www.bcgsc.ca/downloads/morinlab/reference/af-only-gnomad.hg38.vcf.g z). Initial variant calls were filtered with FilterMutectCalls. To ensure high-confidence calls, a stringent second filtering step was applied using bcftools ^[27]^, retaining only variants with a depth (DP) ≥20, allele frequency (AF) ≥0.05, and tumor log odds score (TLOD) ≥6.

Functional annotation of filtered variants was performed using the Ensembl Variant Effect Predictor (VEP) web interface ^[30]^. To normalize the number of somatic variants identified in each sample, we calculated the number of high-quality, uniquely aligned reads using samtools ^[27]^. First, duplicate reads were marked using GATK’s MarkDuplicates tool. Then, the resulting BAM files were indexed and processed with samtools view -c -F 0x904 -q 20, which counts primary alignments with a mapping quality (MAPQ) ≥20, while excluding unmapped reads, secondary alignments, and PCR duplicates. The final number of variants per sample was normalized by dividing the total variant count by the number of these high-quality mapped reads, and multiplied by one million. Normalized variant counts were then used to estimate the somatic mutation burden for each sample.

Prior to statistical comparisons, the distribution of normalized somatic mutational burden values was evaluated for the presence of outliers using the Tukey criterion (values exceeding Q3 + 1.5xIQR). Samples identified as outliers were excluded from subsequent statistical analyses. Statistical comparisons were performed in R using two-sample *t*-tests to evaluate differences between groups stratified by NAS score and PNPLA3 genotype.

### Correlation network analysis

In order to explore potential interactions between microbial taxa, functional signatures, and somatic alterations associated with liver injury and genetically associated risk in MASLD, correlation network analyses were conducted separately for all study groups, stratified by NAS score (NAS≥5 and NAS≤4) and by PNPLA3 genotype (GG and CC/CG). The input dataset included: (i) metacommunity composition according to the read counts assigned at the species level across all liver metagenomes, (ii) the normalized number of somatic variants detected per sample (expressed as variants per million high-quality mapped reads), (iii) InterPro protein families significantly associated with either NAS score or PNPLA3 genotype (*q*<0.01, present in ≥10 samples), and (iv) statistically significant eggNOG orthologous groups under the same inclusion criteria.

Correlation analyses were performed using the Hmisc R package ^[31]^, based on Spearman’s correlation coefficients. Only significant correlations, defined as those with |ρ|>0.75 and *p*<0.01, were retained for network construction. Weighted correlation networks were generated using the R package igraph v.1.2.7. ^[32]^, while Cytoscape v.3.71 ^[33]^ was used for network visualization. To infer the regulatory relevance of each variable or node in the system represented by these networks and identify potential keystones, topological parameters were computed with the igraph package ^[32]^, including degree centrality (*i.e.*, the number of connections each node has in a network) and eigenvector centrality scores (*i.e.*, the extent to which a node is part of the path connecting other nodes, weighting their degree scores). Modularity analyses (*i.e.*, the degree to which a network can be divided into independent groups or modules) were performed using the optimal cluster algorithm when computationally possible, or using Louvain’s clustering algorithm otherwise ^[34,35]^.

### Statistical analyses

Descriptive statistics were expressed as mean with standard deviation (SD), or as absolute numbers with corresponding percentages. Categorical variables were analyzed using contingency tables and the Fisher’s exact test, while continuous variables were assessed with the Mann–Whitney U test. Statistical analyses were conducted using the Statistical Package for the Social Sciences (SPSS), v.18.0 (SPSS Inc., Chicago, IL, USA), considering a *p*-value<0.05 as statistically significant.

Additional statistical analyses were carried out using RStudio (version 1.3.193; RStudio Team, 2020), and graphical outputs were generated with the ggplot2 package ^[36]^.

## RESULTS

### Clinical, epidemiological and genetic data of the recruited cohort

The main characteristics of the 30 recruited subjects is shown in **Table 1**. As shown in this table, both groups displayed a high degree of homogeneity regarding any variable that could be considered a confounding factor, minimizing their potential effect in the analysis of the hepatic metagenome.

**Table 1.**
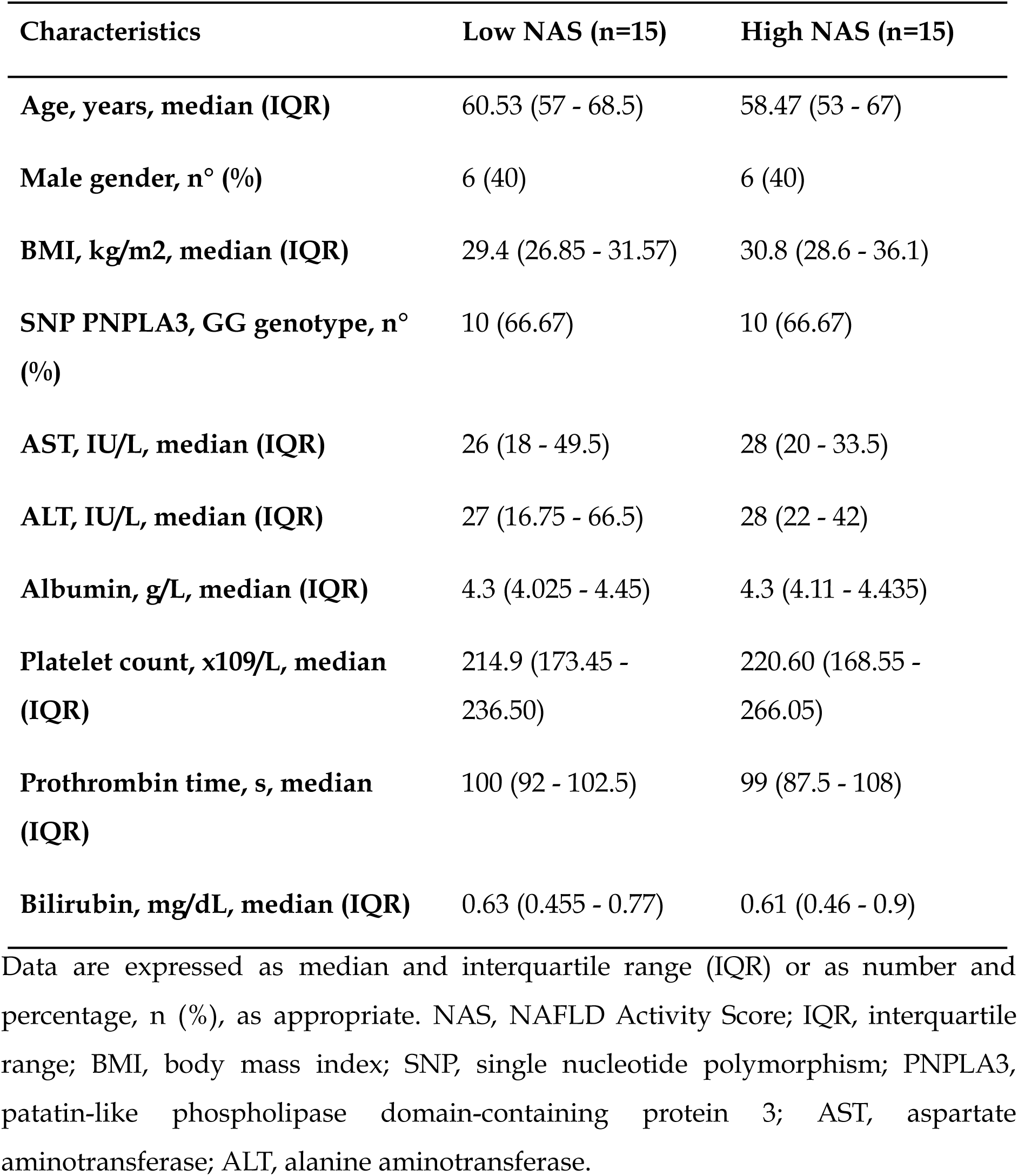
Characteristics of the recruited subjects.

### Liver tissue metagenome profile

First, the liver metagenome profile of each group was evaluated for the proportion that each microorganism domain represented, revealing bacterial genes as the most abundant ones, followed by fungal, protozoan, viral and archaeal (**Table 2**). When the number of reads belonging to each one was compared between groups, bacteria (*p*<1.00E-5), fungi (*p*<1.00E-5), protozoa (*p*<1.00E-5) and archaea (*p*=0.03) were significantly more abundant in patients from the low NAS group (**Table 2**).

**Table 2.** Distribution of metagenomic read counts across microbial domains according to NAS score.

|  | Low NAS group | High NAS group |
| --- | --- | --- |
| **Bacterial reads**<br>(number(%)) | 22276 (3.197%) | 11852 (2.323%)<sup>a</sup> |
| **Fungal reads**<br>(number(%)) | 598 (0.086%) | 249 (0.049%)<sup>a</sup> |
| **Viral reads** (number(%)) | 114 (0.016%) | 61 (0.012%) |
| **Archaeal reads**<br>(number(%)) | 16 (0.002%) | 3 (<0.001%)<sup>a</sup> |
| **Protozoan reads**<br>(number(%)) | 565 (0.081%) | 237 (0.047%)<sup>a</sup> |

Total number of shotgun metagenomic reads assigned to each microbial domain in groups of patients stratified by histological severity, defined as low NAS and high NAS groups. Values are reported as absolute read counts, with the percentage of total reads within each group indicated in parentheses. <sup>a</sup> $P < 0.05$ vs low NAS group when Fisher's exact test with Yates' continuity correction was performed.

The same type of analysis was carried out taking into account the PNPLA3 genotype of the subjects. In this sense, although bacteria dominated the liver microbiome of both groups, the proportion of fungal reads was higher than that of protozoa in the microbiome of the subjects with the GG genotype but lower in the other group (**Table 3**). When statistically comparing the number of reads of each domain of the metagenome considering the genotype of the subjects, a significantly lower abundance of bacteria (*p*<1.00E-5), fungi (*p*=6.68E-4) and protozoa (*p*=0.03) was identified in the group with GG genotype when compared to the other group (**Table 3**).

**Table 3.**
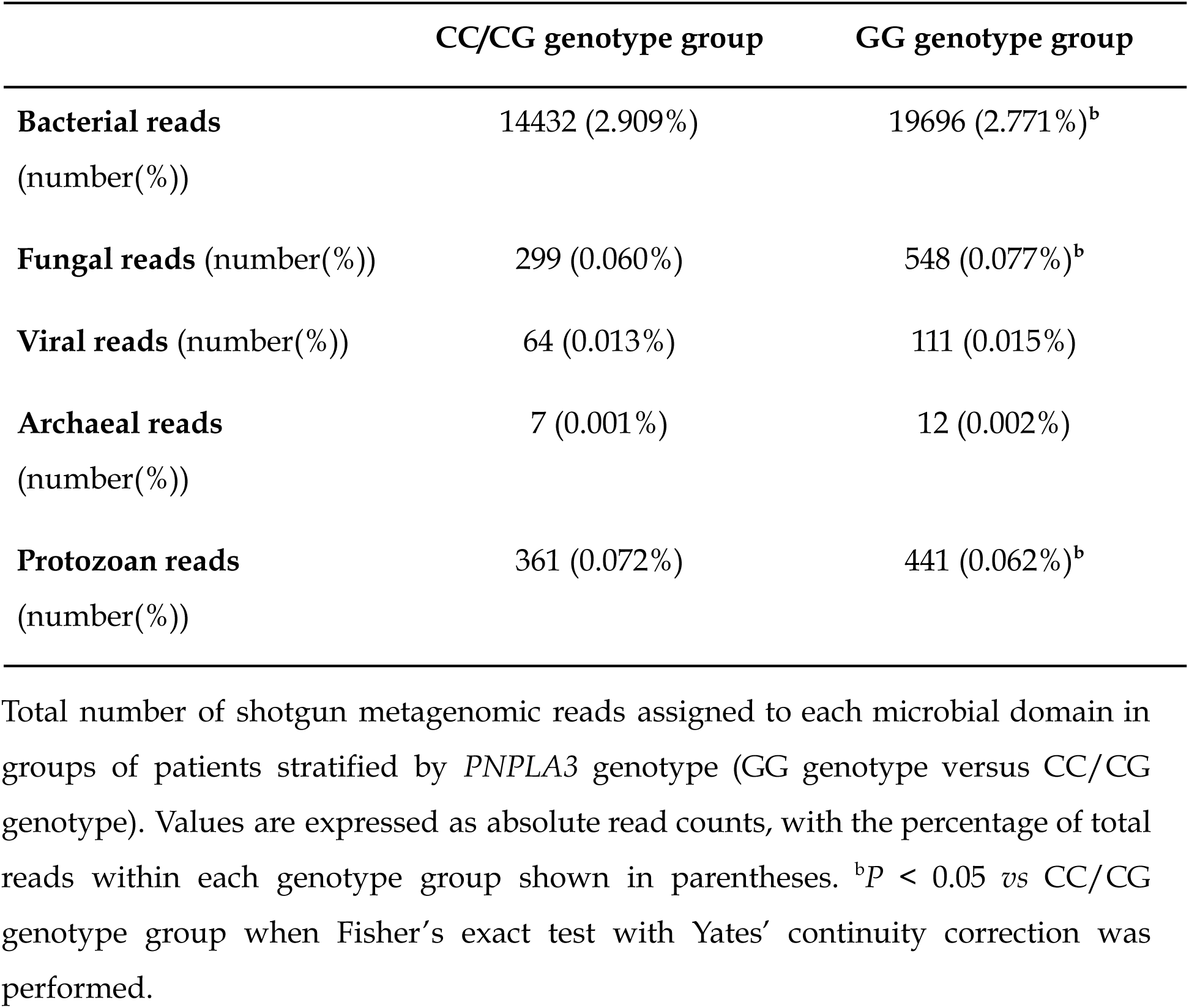
Distribution of metagenomic read counts across microbial domains according to PNPLA3 genotype.

|  | CC/CG genotype group | GG genotype group |
| --- | --- | --- |
| **Bacterial reads**<br>(number(%)) | 14432 (2.909%) | 19696 (2.771%)<sup>b</sup> |
| **Fungal reads** (number(%)) | 299 (0.060%) | 548 (0.077%)<sup>b</sup> |
| **Viral reads** (number(%)) | 64 (0.013%) | 111 (0.015%) |
| **Archaeal reads**<br>(number(%)) | 7 (0.001%) | 12 (0.002%) |
| **Protozoan reads**<br>(number(%)) | 361 (0.072%) | 441 (0.062%)<sup>b</sup> |

Total number of shotgun metagenomic reads assigned to each microbial domain in groups of patients stratified by *PNPLA3* genotype (GG genotype versus CC/CG genotype). Values are expressed as absolute read counts, with the percentage of total reads within each genotype group shown in parentheses. <sup>b</sup> $P < 0.05$ vs CC/CG genotype group when Fisher's exact test with Yates' continuity correction was performed.

Then, metagenome diversity in liver tissue samples was analyzed to evaluate potential differences associated with histological and genetic factors.

No statistically significant differences were observed for alpha diversity indices using Shannon and Chao1 across groups classified by NAS score and PNPLA3 genotype (**Figure 1**).

**Figure 1.**
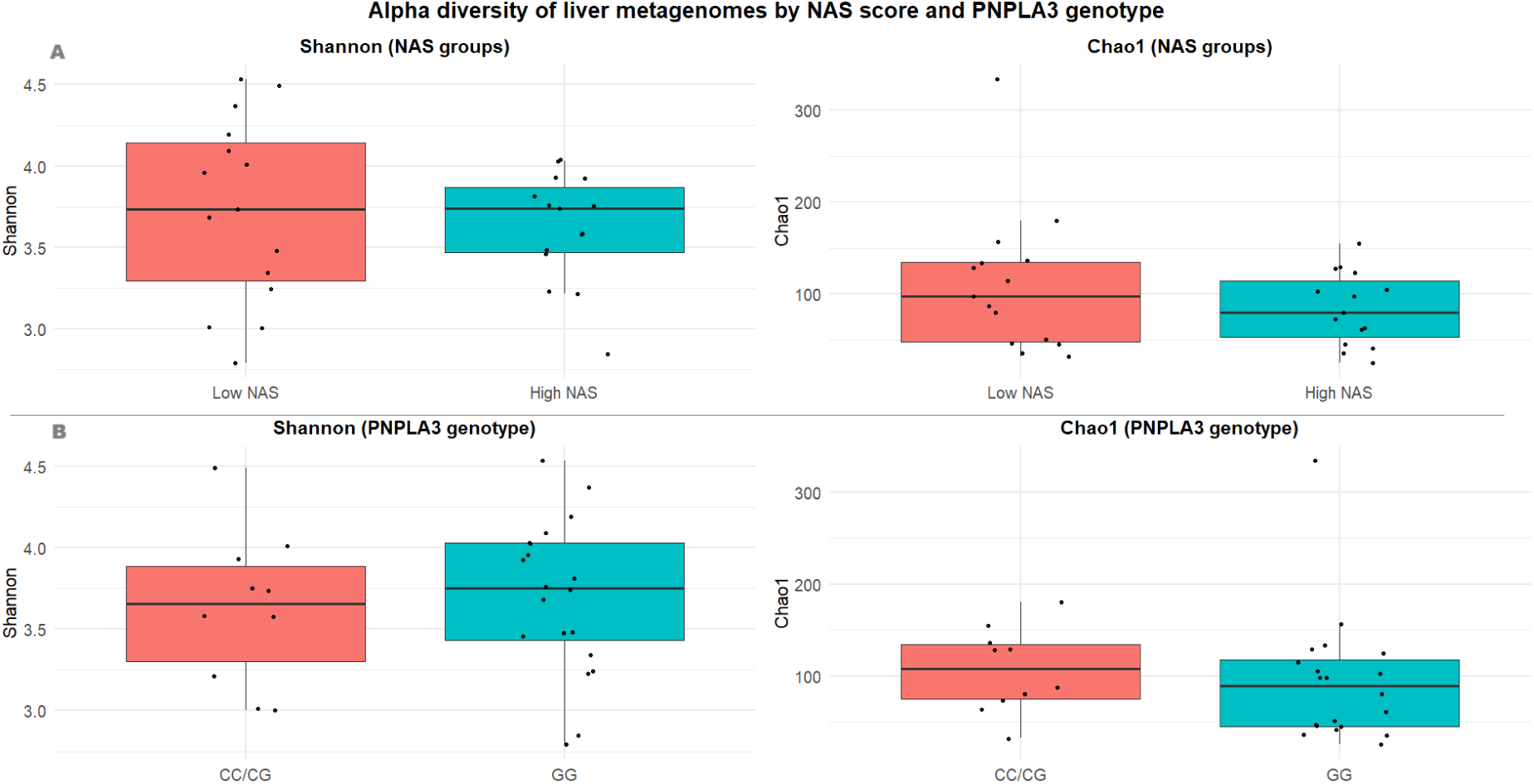
Alpha diversity across MASLD study groups defined by histological severity and PNPLA3 genotype. Boxplots depict Shannon and Chao1 indices at the species level for liver metagenomes stratified by NAS score (A) and PNPLA3 I148M genotype (B). Chao1 values were computed after rarefaction of read counts to the minimum sample sequencing depth using random subsampling without replacement (vegan::rrarefy)^[21]^, whereas Shannon indices were calculated from the original species abundance table. Points represent individual samples; boxplots show the median and interquartile range. Between-group comparisons were performed using the Wilcoxon test, with statistical significance defined as *p*<0.05.

Consistent with the alpha diversity findings, no statistically significant differences in beta diversity were observed between groups for either criterion.

### Compositional and differential abundance analyses

The taxonomic composition of the liver metacommunities was analyzed at the family, genus and species levels. A detailed list of the number of reads assigned to each taxon is provided in **Supplementary Table 1,** while the relative abundance of dominant species identified in these descriptive analyses was plotted in **Figure 2**. Among these dominant microorganisms -defined as those with a relative abundance greater than 0.5% in either group-, bacteria were predominant. However, protozoa (*Plasmodium ovale*), fungi (*Aspergillus fumigatus*, *Malassezia restricta*), viruses (unclassified species of *Siphoviridae, Mycolicibacterium* phage J1), and other eukaryotes (*Trichinella papuae*, *Diploscapter pachys*) were also represented at appreciable relative abundances (**Figure 2**).

**Figure 2.**
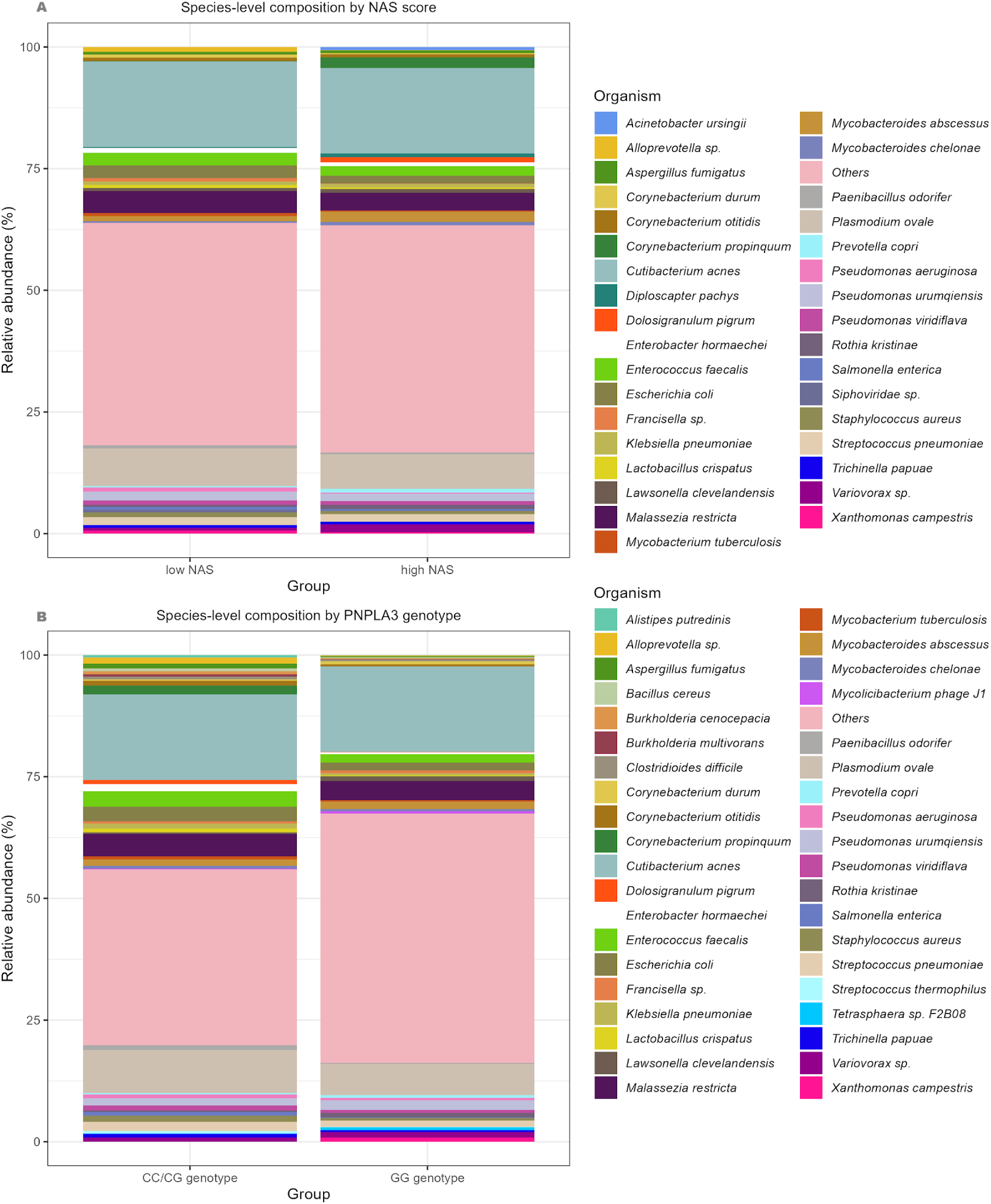
Species-level taxonomic composition of the hepatic metagenome stratified by NAS score and PNPLA3 genotype. Stacked bar plots showing the relative abundance (%) of microbial species detected in liver tissue metagenomes stratified by (A) histological disease severity according to NAS score (low NAS *vs.* high NAS) and by (B) PNPLA3 genotype (CC/CG genotype *vs.* GG risk genotype). Only microbial species present in at least 10 samples and reaching a relative abundance greater than 0.5% in at least one of the compared groups were included in the visualization.

When assessing species prevalence across samples, we found that only 40 of the 1411 identified species were detected in at least one-third of the cohort (**Supplementary Table 1**). This indicates that the metagenome composition of this cohort was highly heterogeneous. Despite marked inter-individual variability in overall community structure, a core set of dominant taxa persisted across disease stages and genetic backgrounds. The identity of the most abundant species was consistent across histological and genotypic strata, with *Cutibacterium acnes* ranking first (low NAS=17.57%, high NAS=17.54%, CC/CG genotype=17.59%, GG genotype=17.53%), followed by *Plasmodium ovale* (low NAS=7.72%, high NAS=7.17%; CC/CG genotype=8.90%, GG genotype=6.38%) and *Malassezia restricta* (low NAS=4.53%, high NAS=3.63%; CC/CG genotype=4.52%, GG genotype=3.95%) (**Figure 2**).

In order to characterize compositional patterns of the liver metagenome in association with the degree of histological damage of the liver in MASLD, a differential abundance analysis was performed at different taxonomic levels. **Supplementary Table 2** provides a comprehensive list of taxa with statistically significant differences between groups, whereas **Figure 3** displays the top 10 taxa with the highest fold-change coefficients at each taxonomic level.

**Figure 3.**
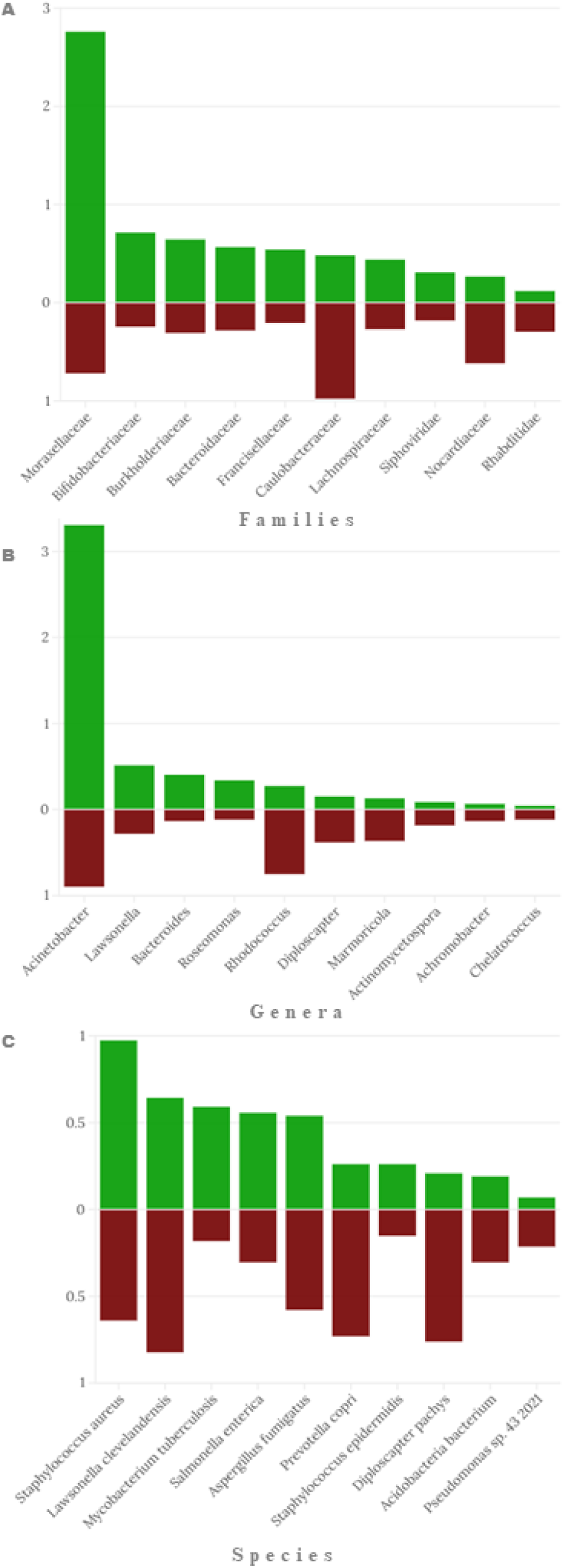
Differential taxonomic abundance of the hepatic microbiome according to histological severity in MASLD. Differential abundance analysis of the liver metagenome comparing patients with low (green) *vs.* high (red) NAS scores. The figure displays relative abundance the top 10 taxa with the highest fold-change coefficients at each taxonomic rank, stratified by histological severity. Panels show results at the family (A), genus (B), and species (C) levels.

Overall, 23 species, 24 genera, and 23 families exhibited significant differences between low and high NAS groups. Although most alterations involved bacterial taxa, fungi (*Aspergillus fumigatus*), viruses (*Mycolicibacterium* phage J1, *Siphoviridae*, *Podoviridae*), and eukaryotes (*Diploscapter pachys*, *Rhabditidae*) were also affected (**Supplementary Table 2**). The most pronounced enrichments in the high NAS group included *Pseudomonas* sp. 43_2001, *Rhodococcus*, *Marmoricola*, *Caulobacteraceae*, and the nematode family *Rhabditidae*, whereas marked depletions were observed for *Staphylococcus epidermidis*, *Acinetobacter*, *Roseomonas*, *Moraxellaceae*, *Bifidobacteraceae*, and *Siphoviridae* (**Figure 3**).

A similar analysis stratified by PNPLA3 genotype revealed substantial compositional differences (**Supplementary Table 3**; **Figure 4**). In total, 22 species, 39 genera, and 36 families were differentially abundant between GG and CC/CG groups, with a predominant shift of depletion of taxa in GG carriers. While bacteria constituted the majority of altered taxa, fungal (*Aspergillus fumigatus*), viral (*Mycolicibacterium* phage J1, *Podoviridae*), and eukaryotic (*Trichinella papuae*, *Diploscapter pachys*, *Rhabditidae*, *Trichinellidae*) lineages were also significantly affected (**Supplementary Table 3**). Among the most notable genotype-associated shifts, *Salmonella enterica* and *Aspergillus fumigatus* were markedly reduced in GG carriers, whereas *Pseudonocardia*, *Propionibacterium*, *Erythrobacteraceae*, and *Oxalobacteraceae* were enriched. In contrast, *Acinetobacter*, *Pantoea*, *Moraxellaceae*, *Pasteurellaceae*, and *Trichinellidae* displayed the strongest depletion in the GG genotype group (**Figure 4**).

**Figure 4.**
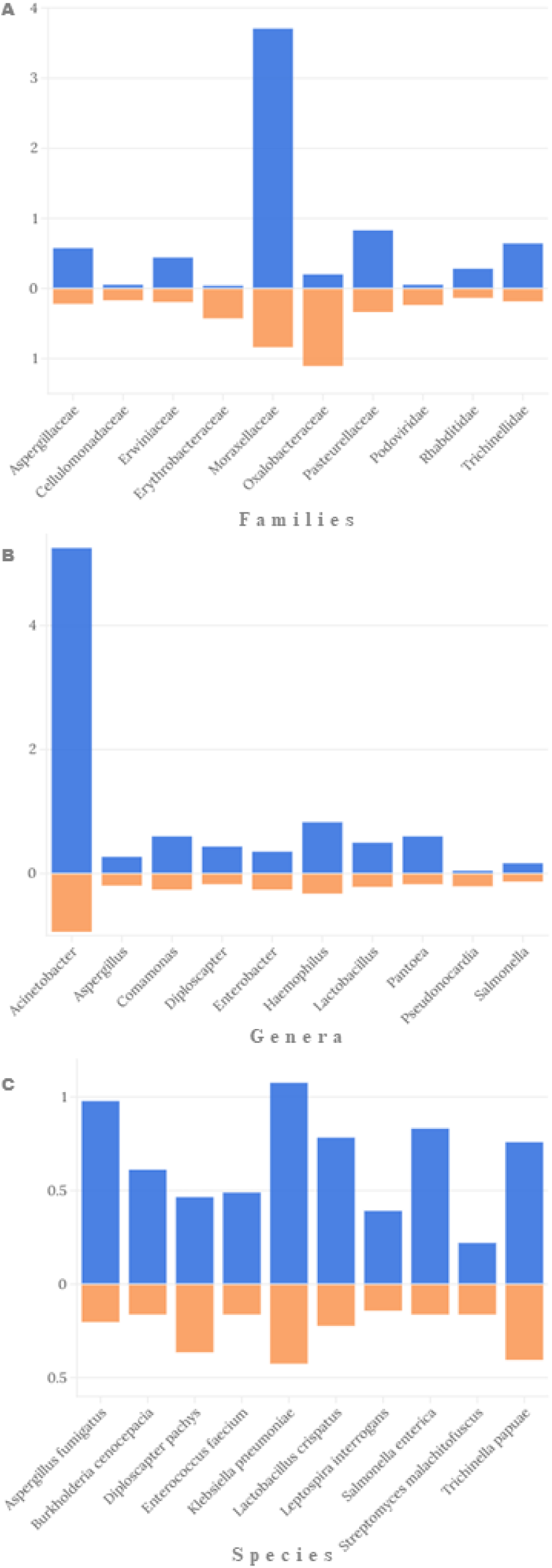
Differential taxonomic abundance of the hepatic microbiome according to *PNPLA3* genotype in MASLD. Differential abundance analysis of the liver metagenome comparing patients carrying the *PNPLA3* CC/CG genotype (blue) *vs.* the GG genotype (orange). The figure displays the relative abundance of the top 10 taxa with the highest fold-change coefficients at each taxonomic rank, stratified by *PNPLA3* genotype. Panels show results at the family (A), genus (B), and species (C) levels.

### Functional enrichment analysis

After evaluating the impact of the degree of histological damage of the liver and of host PNPLA3 genotype on the taxonomic composition of the liver metagenome, we assessed changes in functional gene potential in association with these traits, through a functional enrichment analysis. To achieve a more robust functional characterization, these comparisons were performed considering two different databases: InterPro and eggNOG.

When comparing the functional potential of subjects according to the degree of histological injury, we identified significant differences in the abundance of genes belonging to 22 InterPro protein families (18 enriched and 4 depleted in the high NAS group), and 16 eggNOG ortholog groups (11 enriched and 5 depleted in the high NAS group). Among the strongest changes observed, we found that the high NAS group exhibited a notable increase in the abundance of genes belonging to the TonB-dependent receptor-like protein family (coef.=2.33), the RND efflux pump (coef.=1.91) and the HTH AraC-type transcriptional regulator family (coef.=1.71).

Additionally, a significant enrichment was observed in the eggNOG orthologous groups associated with ABC-type glycerol-3-phosphate transport system (coef.=2.52) and the ATP-dependent Clp protease ATP-binding subunit (coef.=1.70) (**Figure 5**).

**Figure 5.**
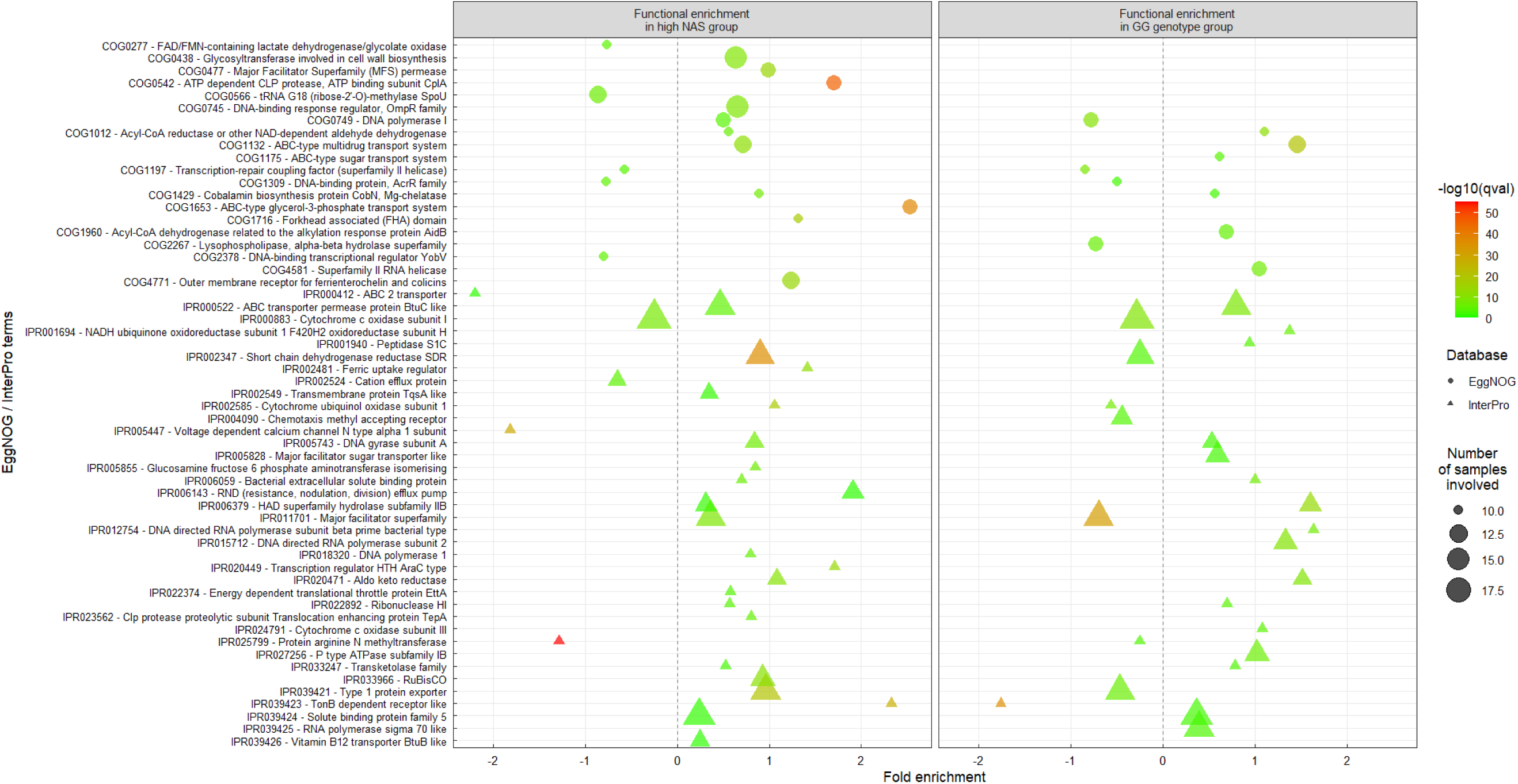
Functional enrichment patterns of the hepatic metagenomes associated with histological severity and *PNPLA3* genotype in MASLD. Enrichment profiles compare subjects with high NAS versus low NAS scores, and carriers of the *PNPLA3* GG genotype versus CC/CG genotypes. Functional annotations were derived from two complementary databases, InterPro and eggNOG. Point color represents the level of statistical significance, expressed as −log10(q-value). Triangular markers correspond to functional features annotated in the InterPro database, whereas circular markers represent eggNOG annotations. Point size reflects the number of samples in which each feature was detected. Only functions observed in over 10 samples and displaying statistically significant enrichment between groups are shown.

When assessing the functional potential of subjects based on PNPLA3 genotype, we identified significant variations in the abundance of genes associated with 24 InterPro protein families (16 enriched and 8 depleted in the risk genotype group) and 10 eggNOG orthologous groups (6 enriched and 4 depleted in the same group). Notably, the GG genotype group displayed a substantial increase in genes linked to the InterPro family DNA-directed RNA polymerase subunit beta-prime, bacterial type (coef.=1.64) and HAD superfamily hydrolase, subfamily IIB (coef.=1.60), as well as the eggNOG orthologous group ABC-type multidrug transport system (coef.=1.46). In contrast, genes belonging to the InterPro family TonB-dependent receptor-like (coef.=-1.76) and the eggNOG orthologous group transcription-repair coupling factor (coef.=-0.84) were notably reduced (**Figure 5**).

### Somatic mutational burden analysis

To evaluate potential associations between somatic mutational burden and disease severity or genetic background, we compared the normalized number of somatic variants across NAS score and PNPLA3 subgroups. A detailed list of somatic mutational burden analysis per sample can be explored at **Supplementary Table 4**.

Two samples were identified as outliers according to the Tukey criterion and were excluded from the statistical analyses. When somatic mutational burden was assessed regarding the disease severity, patients from the high NAS group exhibited a higher mean normalized variant count than those from the low NAS group (148.85 *vs* 132.65); however, this difference was not statistically significant (t=1.92, *p*=0.067). Similarly, when stratifying by PNPLA3 genotype, individuals harboring the risk-associated allele showed a trend toward increased somatic mutational burden compared to non-carriers (144.14 *vs* 134.32), yet the difference did not reach statistical significance (t=-1.08, *p*=0.293).

### Correlation network analysis

In patients with high NAS scores, the resulting network contained 114 nodes connected by 151 significant edges, with an average connectivity (*i.e.*, average node degree) of 2.64 edges per node (**Figure 6**). Among these associations, only one represented a negative correlation, denoting an overall synergistic network structure. Nodes showing the highest degree centrality scores were *Shinella fusca*, *Sphingomonas leidyi*, *Acidobacteria bacterium* RIFCSPLOWO2_12_FULL_67_14b, and *Sphingomonas* sp. 67-41. The eigenvector centrality score identified *Sphingomonas leidyi* as the most influential node, followed by *Shinella fusca*, *Acidobacteria bacterium* RIFCSPLOWO2_12_FULL_67_14b, and *Sphingomonas* sp. 67-41. All of these nodes belonged to the same module. Their strong eigenvector values highlight the strong integration of these taxa within the core of the network, and their collective influence over other nodes, confirming *S. leidyi* as the top-ranking keystone species in terms of overall network prestige. Community structure analysis revealed a 25 clusters partition (modularity score = 0.78), with a single dominant cluster (Module 3) encompassing most of the high-degree and high-prestige nodes. Several smaller peripheral communities exhibiting low centrality were identified, potentially representing specialized or context-dependent associations with minimal contribution to the overall network coherence. Among functional components, the InterPro family IPR005855 (glucosamine-fructose-6-phosphate aminotransferase) displayed high connectivity, suggesting it has a role as a functional hub within the main microbial cluster. Noteworthy, a cross-layer association was identified between *Salmonella enterica* and the somatic variant burden in hepatic tissue, linking specific microbial profiles to genomic instability signals detected in the host.

**Figure 6.**
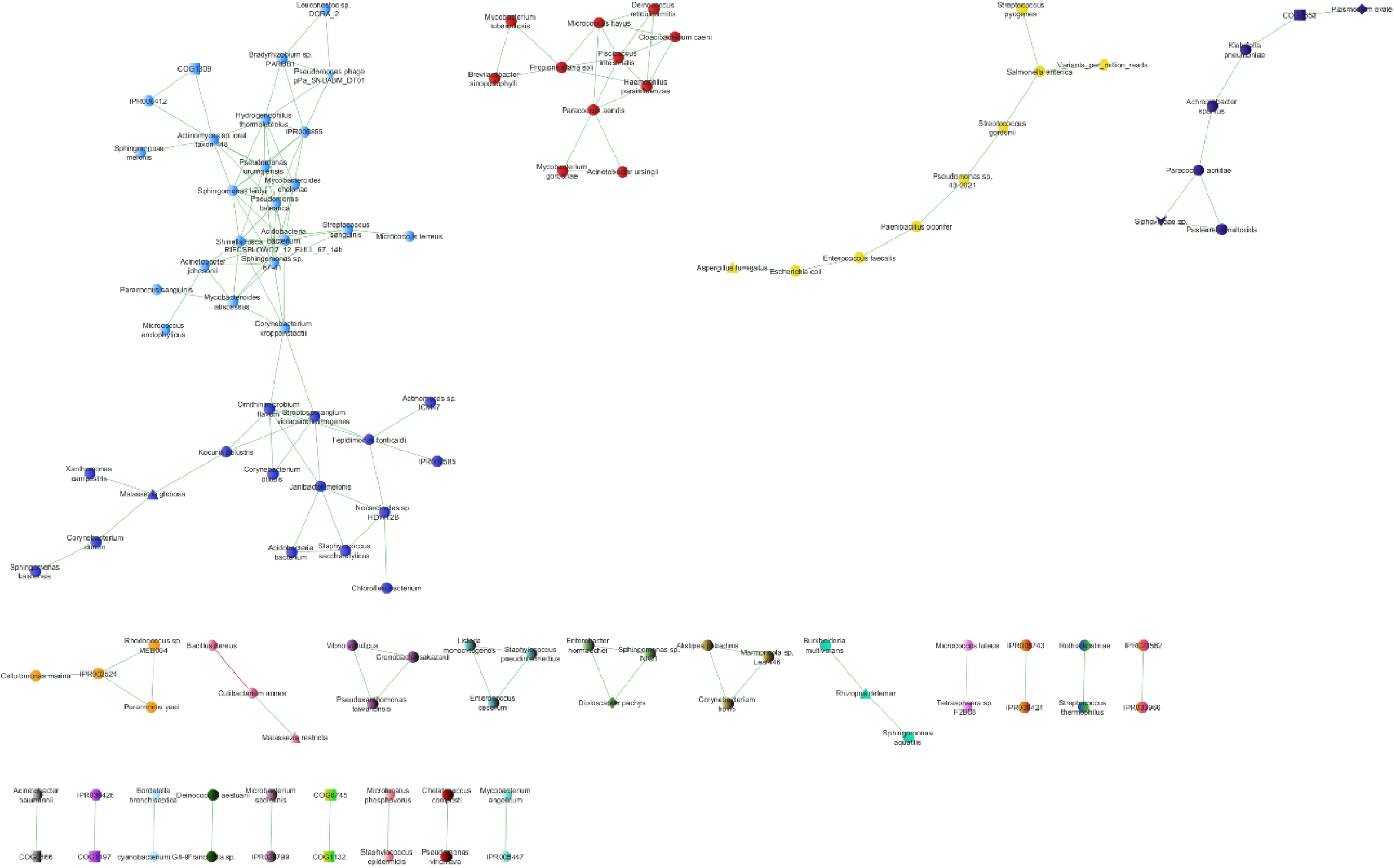
Spearman correlation network of hepatic microbial, functional, and somatic variables in high NAS MASLD patients. Weighted Spearman correlation network representing significant associations among microbial taxa, functional signatures, and somatic mutational burden in patients with high histological activity (NAS ≥5). Networks were constructed using species-level microbial read counts, normalized somatic mutational burden (variants per million high-quality mapped reads), and InterPro and eggNOG functional features significantly associated with NAS score (q<0.01, present in ≥10 samples). Only correlations with |ρ|>0.75 and p<0.01 were retained. Green edges indicate positive correlations, whereas red edges represent negative correlations. Nodes sharing the same color belong to the same network cluster. Node shapes denote variable type: bacteria (circles), viruses (arrowheads), protozoa and eukaryotic parasites (diamonds), fungi (triangles), eggNOG functional annotations (squares), InterPro protein families (octagons), and somatic mutational burden (hexagons).

The low NAS network was larger and more densely connected, comprising 132 nodes and 335 significant edges, resulting in an average connectivity of 5.08, a substantially higher value compared to the high NAS network (**Figure 7**). No negative correlations were observed in this network. Nodes with the highest degree centrality included *Brevibacterium* sp. CS2, the InterPro family IPR020449 (AraC-type transcriptional regulator), *Cutibacterium avidum* and *Hymenobacter* sp. Eigenvector centrality highlighted *Cutibacterium avidum* as the most influential node. Due to the elevated number of nodes and edges in this network, the application of the cluster optimal community detection algorithm was computationally infeasible for this network, as it did not converge under the available resources. Consequently, community structure was inferred using Louvain’s algorithm, which provides an efficient and scalable approximation of modular organization in large and highly connected networks ^[37]^. The low NAS network displayed 15 clusters (modularity score = 0.62), and no significant correlations with somatic variant burden were detected.

**Figure 7.**
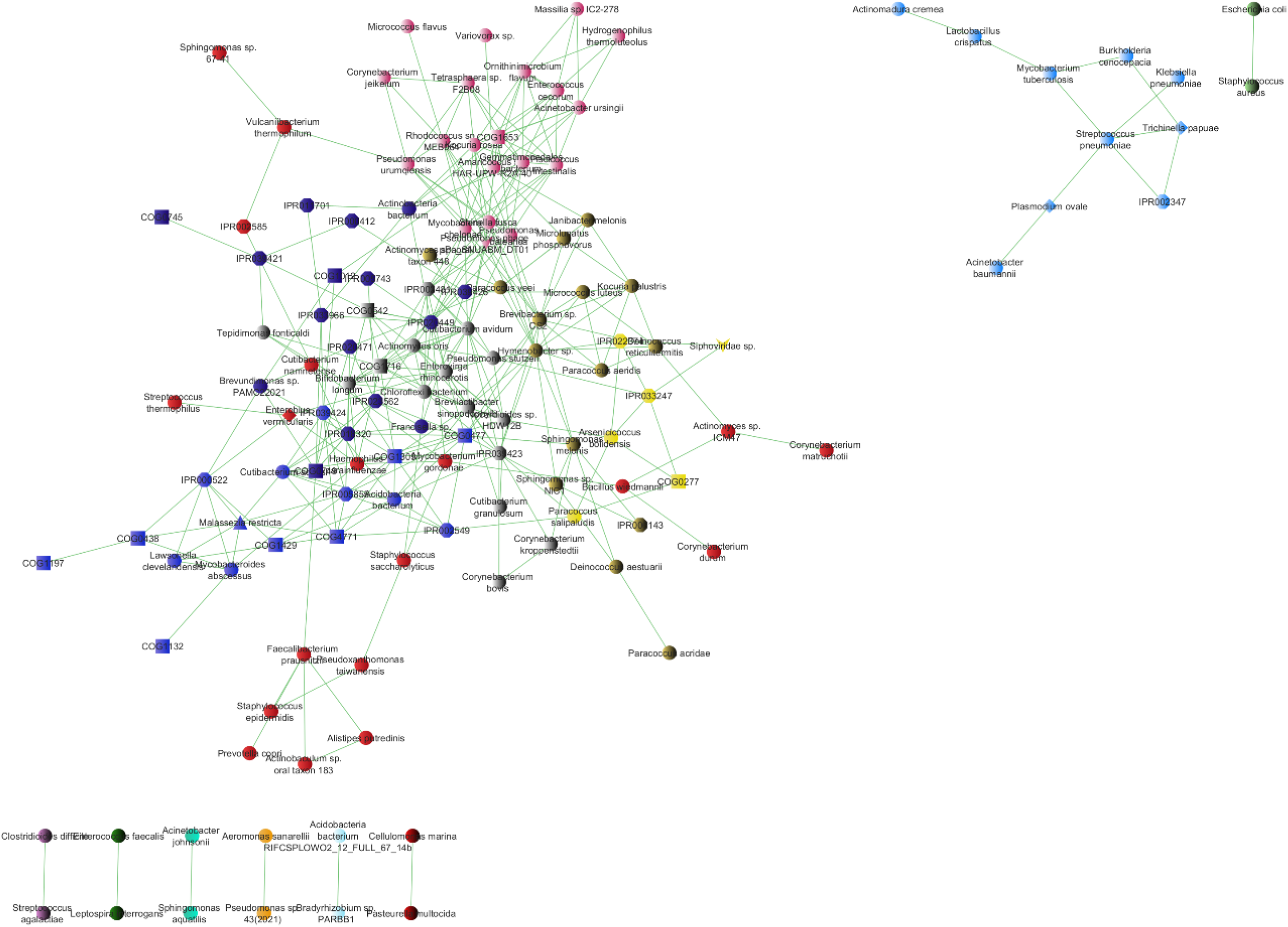
Spearman correlation network of hepatic microbial, functional, and somatic variables in low NAS MASLD patients. Weighted Spearman correlation network depicting significant associations among microbial taxa, functional features, and somatic mutational burden in patients with low histological activity (NAS ≤4). Networks were generated using species-level microbial composition, normalized somatic variant counts, and InterPro and eggNOG functional annotations meeting predefined significance and prevalence criteria (q<0.01, present in ≥10 samples). Only correlations exceeding |ρ|>0.75 with p<0.01 were included. Positive correlations are shown in green and negative correlations in red. Nodes of the same color indicate cluster membership. Node shapes represent variable categories: bacteria (circles), viruses (arrowheads), protozoa and eukaryotic parasites (diamonds), fungi (triangles), eggNOG annotations (squares), InterPro features (octagons), and somatic mutational burden (hexagons).

Stratification by PNPLA3 genotype further revealed distinct network configurations. The network corresponding to PNPLA3 GG carriers consisted of 79 nodes and 107 significant edges, yielding the lowest average connectivity among all groups (1.71) (**Figure 8**). All associations in this network were positive. Nodes with the highest degree centrality included *Hydrogenophilus thermoluteolus*, *Acinetobacter ursingii*, *Brevibacterium* sp. CS2, and *Sphingomonas* sp. 67-41, while eigenvector centrality consistently identified *Hydrogenophilus thermoluteolus* and *Acinetobacter ursingii*as the most influential node. The network was partitioned into 16 clusters, with a modularity score of 0.71.

**Figure 8.**
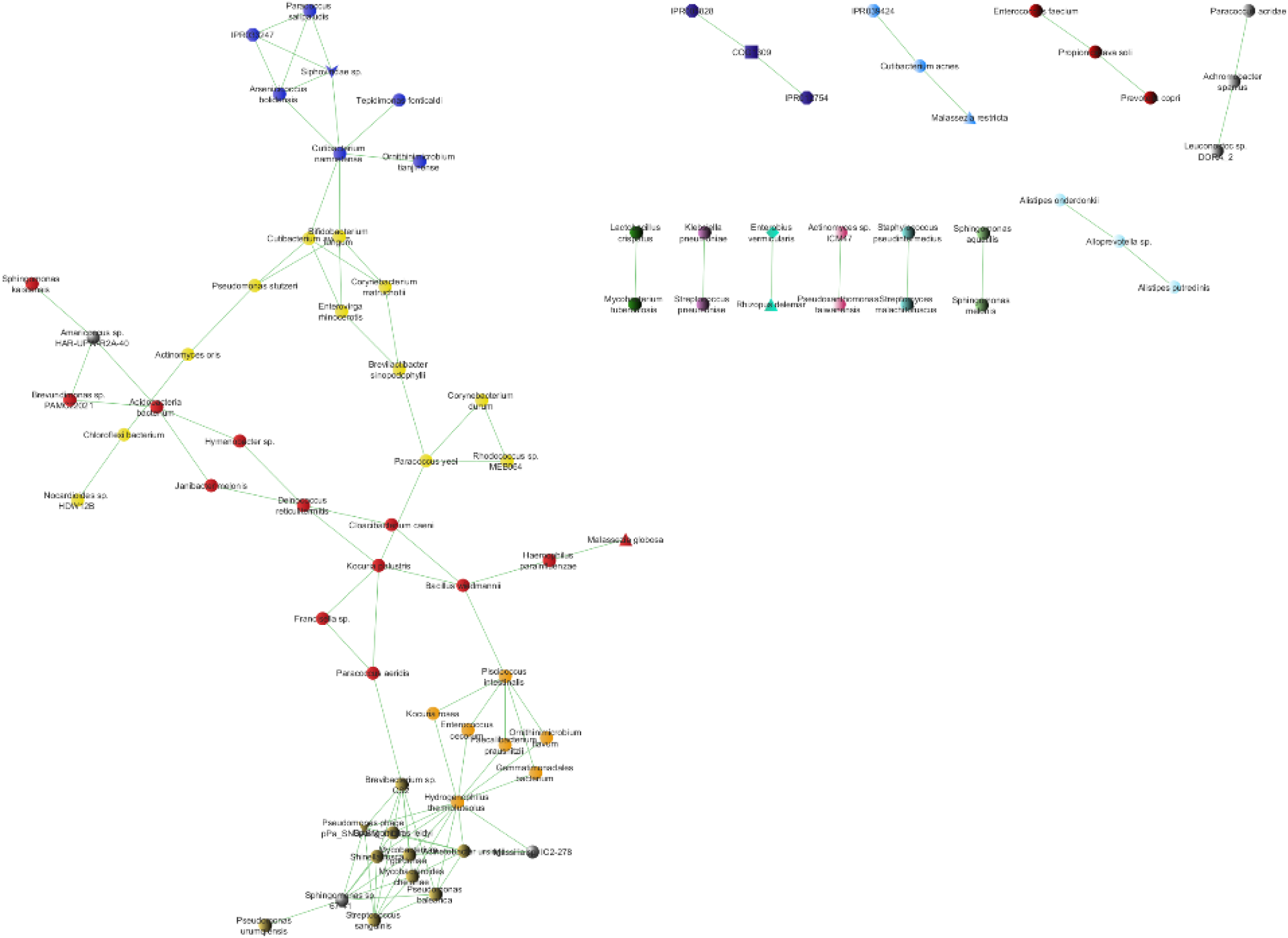
Spearman correlation network of hepatic microbial, functional, and somatic variables in *PNPLA3* GG genotype carriers. Weighted Spearman correlation network illustrating significant associations among microbial taxa, functional signatures, and somatic mutational burden in subjects carrying the *PNPLA3* GG genotype. Networks were constructed using species-level microbial read counts, normalized somatic variant burden, and InterPro and eggNOG features significantly associated with *PNPLA3* genotype (q<0.01, present in ≥10 samples). Only correlations with |ρ|>0.75 and p<0.01 were retained. Green edges denote positive correlations and red edges denote negative correlations. Nodes sharing the same color belong to the same cluster. Node shapes indicate variable type: bacteria (circles), viruses (arrowheads), protozoa and eukaryotic parasites (diamonds), fungi (triangles), eggNOG functional annotations (squares), InterPro protein families (octagons), and somatic mutational burden (hexagons).

In contrast, the PNPLA3 CC/CG genotype network was the largest and most heterogeneous, comprising 149 nodes and 288 significant edges, with an average connectivity of 3.87 (**Figure 9**). This network exhibited 21 negative correlations, representing the highest number of antagonistic associations among all groups. Nodes with the highest degree centrality included *Corynebacterium kroppenstedtii*, *Mycobacteroides chelonae*, *Paracoccus sanguinis*, *Pseudomonas balearica*, and *Pseudomonas urumqiensis*, together with the functional node COG1309 (AcrR family DNA-binding protein). Eigenvector centrality revealed a tightly interconnected set of nodes concentrated within a dominant cluster, where *Mycobacteroides chelonae*, *Pseudomonas balearica* and *Pseudomonas urumqiensis* exhibited the highest prestige. This network was partitioned into 20 clusters and exhibited the highest modularity score (0.80). A significant cross-layer association was detected between *Streptococcus gordonii* and hepatic somatic variant burden.

**Figure 9.**
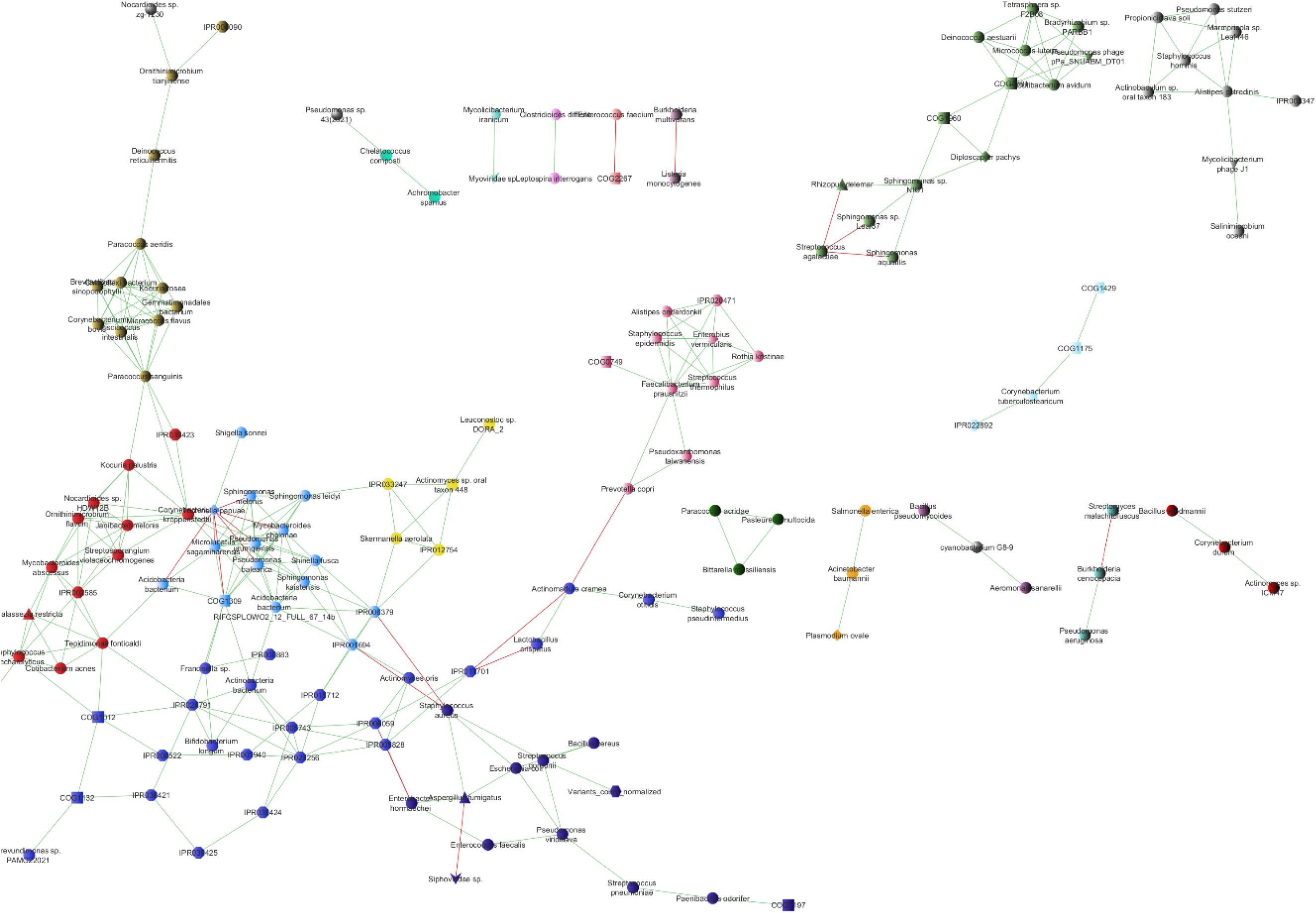
Spearman correlation network of hepatic microbial, functional, and somatic variables in *PNPLA3* CC/CG genotype carriers. Weighted Spearman correlation network depicting significant associations among microbial taxa, functional features, and somatic mutational burden in subjects carrying the *PNPLA3* CC/CG genotype. Networks were generated using species-level microbial composition, normalized somatic variant counts, and InterPro and eggNOG functional annotations ulfilling predefined statistical and prevalence thresholds (q<0.01, present in ≥10 samples). Only correlations with |ρ|>0.75 and p<0.01 were included. Positive correlations are shown in green and negative correlations in red. Nodes of the same color indicate cluster membership. Node shapes reflect variable categories: bacteria (circles), viruses (arrowheads), protozoa and eukaryotic parasites (diamonds), fungi (triangles), eggNOG annotations (squares), InterPro features (octagons), and somatic mutational burden (hexagons).

## DISCUSSION

Most research on MASLD pathophysiology has primarily focused on gut bacterial dysbiosis ^[38]^. However, our findings focus on the hepatic niche, underscoring the importance of non-bacterial components as integral players in the hepatic microbiome ecosystem. Comparative analyses revealed that the relative abundances of bacteria, fungi, protozoa, and archaea were significantly higher in patients with lower histological injury. Similarly, patients with the CC/CG genotype exhibited significantly higher abundances of bacteria, fungi, and protozoa compared to GG genotype carriers. These results support the notion that disease progression and genetic susceptibility in MASLD may be associated with a progressive depletion of microbial abundance, possibly reflecting a more hostile hepatic microenvironment characterized by inflammation, oxidative stress, or altered metabolic composition. The observed reduction in multiple microbial domains in high NAS and GG genotype patients aligns with prior studies describing a loss of taxonomic and functional diversity in the gut microbiome of patients with advanced fibrosis or MASH ^[39]^, as well as decreased fungal diversity in cirrhotic individuals ^[40]^. While the causal direction of this relationship remains unclear, it is plausible that lower microbial colonization in the liver in the above mentioned cases may result from a more restricted microbial translocation through the intestinal barrier or, most likely, an enhanced hepatic immune activation impairing microbial survival.

When microbiome structure was analyzed, no significant differences were found in alpha diversity indices nor in beta diversity between patient groups. These results indicate that although specific microbial domains and taxa vary in abundance, the overall richness and diversity distribution within and between individuals remain relatively stable. This apparent paradox has been previously reported in MASLD-related gut microbiome studies ^[41]^, where microbial shifts associated with disease were driven by specific taxa or functional pathways, rather than global diversity measures. Hence, our results reinforce the idea that microbial community structure alone may not capture functionally relevant dysbiosis, and highlight the need for integrative analyses to identify mechanistic links between microbial activity along the gut-liver axis and disease progression.

Taxonomic profiling of the hepatic metagenome revealed a highly diverse yet sparsely distributed microbial landscape, with most species detected in only a handful of samples. Out of the 1411 species identified, only 40 appeared in over a third of the 30 samples. This limited and highly individualized presence aligns with the concept of a low-biomass hepatic microbiome shaped by selective pressures ^[42]^. It likely reflects the restricted microbial translocation that occurs across the gut-liver axis, where the portal vein primarily carries microbial byproducts like lipopolysaccharides and metabolites rather than live organisms ^[9,43]^. Once in the liver, these components are swiftly recognized and neutralized by resident immune cells, such as Kupffer cells and liver sinusoidal endothelial cells, which help preserve hepatic immune tolerance and prevent microbial colonization. Moreover, the liver itself plays an active role in shaping intestinal microbial communities through bile acid secretion and IgA production, forming part of a bidirectional feedback system essential to gut-liver homeostasis ^[44]^. Altogether, these highly complex mechanisms likely contribute to the low ubiquity of most taxa observed in the hepatic microbial communities of patients with MASLD.

Among the most prevalent microbial taxa detected in liver samples, several non-bacterial organisms consistently appeared across comparisons based on NAS scores and PNPLA3 genotype. These included eukaryotes (fungi and protozoa) as well as viruses. These findings reinforce the polymicrobial nature of the hepatic microbiome and suggest that non-bacterial taxa may have underappreciated roles in the pathophysiology of MASLD. Fungal species like *Malassezia restricta* and *Aspergillus fumigatus* have previously been associated with chronic inflammation and tissue fibrosis. Notably, *Malassezia* spp., which are thought to exacerbate inflammatory liver conditions through immune activation and interaction with bile ducts, have been detected in the gut and liver of patients with hepatocellular carcinoma and primary sclerosing cholangitis ^[45,46]^.

Likewise, the nematodes *Trichinella papuae* and *Diploscapter pachys* were the most abundant eukaryotic taxa, and also emerged as differentially abundant in both histological severity- and genotype-based comparisons. While the detection of helminth-derived sequences in hepatic samples may partially reflect environmental exposure or dietary sources, the consistent presence of these taxa in multiple individuals raises the possibility that they contribute functionally to the hepatic microenvironment. Of note, *D. pachys* is known to produce antifungal peptides targeting filamentous fungi ^[47]^, suggesting a possible role in modulating fungal colonization or immune tone. These taxa could be interpreted within the framework of pathobionts, *i.e.* organisms that are typically commensal but may become pathogenic in the context of host immune dysfunction or ecological imbalance. This concept is increasingly recognized in mucosal microbiology, where context-dependent virulence plays a central role in shaping disease outcomes ^[48]^.

Viral sequences, particularly those belonging to the Siphoviridae family and the species *Mycolicibacterium* phage J1, were also abundant in the liver metagenomes of MASLD patients. Many recent findings highlight the virome as a critical and underexplored regulator of host-microbiota interactions ^[49]^. Bacteriophages, such as those from the Siphoviridae family, are known to shape bacterial community structure and function, and may modulate immune responses indirectly by influencing bacterial antigen availability, lysogeny, or lysis-driven endotoxin release ^[50]^. In the context of MASLD, such phage-bacteria dynamics may contribute to the observed enrichment of LPS-rich Enterobacteria (*e.g. Salmonella enterica*) in patients with high NAS scores.

To better understand the ecological dynamics of the hepatic microbiome in MASLD progression, we conducted a differential abundance analysis. Several of the differentially abundant taxa identified in this study have been previously associated with metabolic and immune regulatory processes. *Staphylococcus epidermidis*, which was significantly reduced in the high NAS group, is a commensal skin and mucosal bacterium known for its immunomodulatory properties, including the regulation of TLR signaling and the promotion of epithelial barrier integrity. Its depletion has been linked to increased susceptibility to inflammatory conditions ^[51,52]^.

Conversely, the increased abundance of *Pseudomonas sp*. in the high NAS group may be of relevance, as certain *Pseudomonas* species can significantly modulate inflammatory responses in various tissues, primarily through cytokine production and quorum-sensing pathways, highlighting their role in exacerbating inflammatory conditions ^[53,54]^. These taxa are potent activators of TLR4 signaling in hepatocytes and Kupffer cells, promoting inflammation, glutaminase-1 expression, and hyperammonemia, processes implicated in the transition from MASLD to MASH ^[41]^. These findings echo prior reports identifying a *Proteobacteria*-dominated hepatic signature in obese individuals, and associate this shift with metabolic inflammation and liver injury, particularly in non-morbidly obese MASLD patients ^[55,56]^.

Similarly, the enrichment of the *Rhodococcus* and *Marmoricola* genera within the Actinobacteria phylum could indicate shifts in lipid metabolism and immune activation, given their known capacity for hydrocarbon degradation and interaction with host lipid pathways ^[57,58]^.

At the viral level, the depletion of the *Siphoviridae* family was consistently observed among patients with advanced histological injury, and carriers of the PNPLA3 risk genotype. Such contraction may have immunological consequences, as some bacteriophages have been proposed to play a role in shaping the gut and hepatic microbiota through bacterial predation, potentially influencing host immune homeostasis. A reduction in these viral populations could reflect broader dysbiosis associated with hepatic injury ^[59,60]^. Moreover, this reduced abundance of this viral family in the liver of patients with advanced histological injury, and carriers of the PNPLA3 risk genotype is in line with our previous gut metatranscriptomic analysis, as this family was also found to be transcriptionally less active in the gut of MASLD patients compared to a healthy cohort, and in that of MASH patients compared to those with simple steatosis ^[13]^. These observations could be understood under the hypothesis that *Siphoviridae* phages are functionally depleted in the intestinal ecosystem under more severe disease states, which may limit their capacity to persist, translocate, or be selectively seeded into the hepatic niche ^[9]^. This reproducible signal across distinct biological compartments and methodological approaches highlights the potential centrality of *Siphoviridae* in MASLD-associated microbial dysbiosis. Given that bacteriophages of this family are thought to regulate bacterial populations through predation, their reduction may disrupt bacterial community balance, favoring the expansion of pro-inflammatory taxa and thereby shaping host immune responses ^[50]^. Such alterations could contribute to the immunometabolic disturbances characteristic of MASLD, linking phage dynamics to both microbial ecology and host pathophysiology.

Notably, many species commonly considered as pathobionts were found to be significantly depleted among carriers of the PNPLA3 risk allele, including *Aspergillus fumigatus*, *Enterococcus faecium*, *Klebsiella pneumoniae*, *Leptospira interrogans*, *Salmonella enterica*, or *Trichinella papuae*. Although this result may be paradoxical at first sight, it aligns with the idea that the pathogenic potential of pathobionts is context-dependent ^[48,61]^. Jochum *et al*. highlight that microorganisms classified as pathobionts can also exert beneficial or homeostatic functions depending on host immunity and microbiota configuration. From this perspective, the decreased abundance of pathobionts in PNPLA3 GG carriers could reflect a genotype-driven shift in the gut-liver ecosystem toward a less inflammatory and more restrictive barrier phenotype. Indeed, our previous integrative analysis of the intestinal transcriptome and metabolome in MASLD showed that the PNPLA3 GG genotype was associated with a downregulation of TLR signaling pathways, together with a suppression of innate immune activation and phage-mediated inflammatory responses ^[13]^. The PNPLA3 risk allele is known to alter hepatic lipid metabolism and immune tone, potentially influencing microbial colonization through changes in bile acid profiles, antimicrobial peptide secretion, or immune surveillance mechanisms ^[12,62]^. Consequently, the observed depletion may not necessarily indicate a loss of microbial diversity but rather a form of selective ecological equilibrium, where the metabolic and immunological environment shaped by PNPLA3 constrains the pathogenic effects of pathobionts.

The most pronounced alterations associated with MASLD progression and PNPLA3 genotype are primarily functional rather than taxonomic, as evidenced by analysis of the fold-change coefficients. This suggests that the adaptive capacity of the hepatic microbial ecosystem depends more on metabolic versatility than on compositional shifts. In patients with high NAS scores, an enrichment of genes encoding TonB-dependent receptor-like proteins (IPR039423) and ferric uptake regulators (IPR002481) was observed, consistent with an increased microbial demand for iron acquisition and transport. Iron overload is a recognized feature of advanced MASLD, contributing to oxidative stress and hepatocellular injury ^[63,64]^. TonB systems are critical virulence determinants that enable bacteria to thrive in iron-rich microenvironments by facilitating the uptake of siderophore-bound ferric ions and corrinoids, including vitamin B12, which may explain their selective expansion under pro-oxidative hepatic conditions ^[65,66]^. Regarding these TonB systems, the inverse pattern was detected in carriers of the PNPLA3 GG genotype, which suggests that the pathogenic mechanisms underlying MASLD in this genetic background may involve pathways independent of hepatic iron metabolism. Indeed, PNPLA3-driven steatosis is known to be mediated through altered lipid remodeling rather than dysmetabolic iron accumulation ^[67]^.

Furthermore, the predominance of genes related to nutrient transport and energy metabolism in both NAS- and PNPLA3-based comparisons points to a functional adaptation of the liver microbiota to nutrient limitation and oxidative stress as fibrosis progresses. These findings are consistent with the “metabolic stress adaptation” paradigm described in the gut–liver axis, where dysbiosis favors microorganisms capable of scavenging scarce substrates and resisting bile acid and ROS-mediated toxicity ^[10]^. Conversely, the depletion of transcriptional and translational genes in high-NAS livers may reflect the loss of metabolically less adaptable species, leading to a reduced biosynthetic activity within the hepatic microbiome ^[68]^. These results support the view that disease progression in MASLD is accompanied by a metabolic reprogramming of the hepatic microbiota rather than by major taxonomic shifts, and that PNPLA3-associated disease involves distinct pathogenesis pathways and exerts specific ecological pressures.

Although our analysis did not reveal statistically significant differences in somatic mutational burden between histological or genetic subgroups, certain trends may suggest underlying biological relevance. Specifically, we observed a higher average burden in patients with greater histological severity and in carriers of the PNPLA3 risk allele, which is consistent with the concept that local genomic instability and microenvironmental stress intensify as metabolic liver injury progresses and in genetically susceptible individuals ^[69]^.

These observations fit within a growing body of evidence indicating that chronic liver disease is accompanied by substantial somatic diversification of hepatocytes. Brunner et al. ^[70]^ showed that cirrhotic livers, including cases of NAFLD-related cirrhosis, harbour a significantly higher mutational burden than normal livers. Building on this, Ng et al. ^[71]^ identified recurrent somatic mutations in metabolic regulators (*e.g*., FOXO1, CIDEB, GPAM) across liver biopsies obtained from patients with chronic liver disease, including NAFLD. Such findings point toward a model where hepatocytes undergo a form of clonal selection, driven by the harsh environment of chronic injury and lipotoxic stress to develop metabolic advantages.

At the malignant end of the spectrum, extremely high tumour mutational burden has been documented only in rare cases of hepatocellular carcinoma, underscoring that large shifts in mutation load are most easily detected in advanced disease and neoplastic contexts ^[72]^.

In contrast, the absence of statistically significant differences in mutational burden is therefore likely multifactorial. First, our cohort represents only pre-cirrhotic MASLD, and we compared subgroups within the same disease rather than healthy versus cirrhotic or cancerous liver. Thus, the expected effect size between these intra-MASLD strata is smaller than between normal and cirrhotic or tumour tissue. Second, the sample size, particularly in the PNPLA3 non-risk group (n = 10), limits statistical power; and the use of low-input FFPE material, together with stringent filtering to minimise artefacts, probably reduces sensitivity for low variant allele frequency events and small clones. Moreover, our estimates of mutational burden are derived at the whole-biopsy level, whereas clone-resolved whole-genome sequencing approaches can detect expansion of individual mutant lineages that may be diluted in bulk tissue measurements. These factors likely explain why observed trends did not reach significance. Future studies leveraging larger cohorts, deeper sequencing, and clonal-resolution approaches will be necessary to clarify the role of somatic mutational load in MASLD progression.

Analysis of the correlation networks revealed that disease stage and host genetic background jointly shape microbial interaction patterns within the liver tissue microenvironment, as marked differences across groups emerged in overall network size, edge density, and connectivity. Conceptually, correlation networks capture statistically robust patterns of co-variation (positive edges) and mutual exclusion (negative edges), which may reflect combinations of shared habitat filtering, metabolic coupling, and/or antagonistic interactions rather than direct causality ^[73]^.

Across NAS-stratified networks, histological progression was accompanied by marked topological simplification. The low NAS group exhibited larger, more connected networks, consistent with a more heterogeneous and permissive hepatic environment in early MASLD. In contrast, high NAS was associated with network contraction, likely reflecting stronger environmental filtering driven by inflammation, oxidative stress, altered bile acid signaling, and metabolic constraints that reduce ecological heterogeneity and limit stable microbial interactions [73–75].

Genotype stratification revealed even greater divergence. The CC/CG network showed higher connectivity, modularity, and uniquely contained multiple negative correlations, features typically associated with niche differentiation and competitive dynamics in complex ecosystems ^[68]^. This could reflect that, in the absence of the risk variant, MASLD progression may involve a more polyfactorial microbial–host interplay in which competitive and mutually exclusive relationships remain detectable ^[74]^.

In contrast, the PNPLA3 GG network was consistently the smallest and least connected, indicating a constrained topology. Given the role of the I148M variant in hepatocellular lipid remodeling, genotype-driven metabolic determinism may reduce hepatic environmental heterogeneity and narrow the range of microbial interactions, partially overriding broader ecological dynamics ^[6]^. This interpretation is supported by emerging evidence that PNPLA3 influences the spatial and metabolic landscape of the liver and shapes microbiome-associated functional signals in MASLD ^[75]^.

Within the high NAS network, a structurally cohesive cluster composed of *Sphingomonas leidyi*, *Sphingomonas* sp. 67-41, *Shinella fusca*, and *Acidobacteria bacterium RIFCSPLOWO2_12_FULL_67_14b* exhibited the highest degree and eigenvector centrality scores, identifying them as keystone nodes. In microbial network theory, such nodes exert disproportionate influence on ecosystem structure and stability ^[73,76]^. Their co-localization within a high-connectivity module supports the existence of a cooperative functional unit in advanced MASLD ^[73]^. Functionally, this module was associated with the InterPro family IPR005855 encoding glucosamine-fructose-6-phosphate aminotransferase (GFAT), the rate-limiting enzyme of the hexosamine biosynthetic pathway. Increased GFAT activity enhances UDP-N-acetylglucosamine production, a precursor for bacterial peptidoglycan synthesis. Enhanced peptidoglycan turnover and muropeptide release may activate intracellular pattern-recognition receptors such as NOD1 and NOD2 in hepatocytes and liver-resident immune cells, pathways implicated in hepatic inflammation, insulin resistance, and steatohepatitis ^[77–79]^. Thus, this main module may reflect either microbial adaptation to a stressed hepatic microenvironment or active participation in amplifying metabolic dysfunction and inflammation, including mechanisms attributed to bacterial sphingolipids in *Sphingomonas* species ^[80]^.

Finally, integration of network topology with somatic mutational burden revealed genotype- and severity-specific associations between microbial signals and genomic instability. In the high NAS network, *Salmonella enterica* gene abundance positively correlated with somatic variant burden. Experimental evidence indicates that genotoxin-producing *S. enterica* can induce tissue-specific DNA damage responses, with oxidative stress–mediated DNA injury predominating in the liver ^[81]^. This provides biological plausibility for the observed association and supports the notion that microbial components with genotoxic or pro-oxidant potential may contribute to genomic instability in MASLD ^[82]^. In the CC/CG network, a similar positive association was observed between *Streptococcus gordonii* abundance and mutational burden. Although classically considered a commensal organism, *S. gordonii* can activate innate immune pathways through its cell wall components and adhesins, promoting low-grade inflammation ^[83]^. Chronic exposure to such inflammatory stimuli may foster a pro-oxidative hepatic microenvironment characterized by increased reactive oxygen and nitrogen species, impaired redox balance, and sustained immune activation — well-recognized drivers of DNA damage accumulation ^[84]^. In this context, the association likely reflects an indirect mechanism whereby immune-mediated oxidative stress creates permissive conditions for mutagenesis rather than a direct genotoxic effect.

When interpreting these results, some limitations of this study must be considered. Among them, one of the main concerns in tissue-based microbiome research involves the use of FFPE specimens, since the preservation process raises the possibility that part of the detected microbial DNA could originate from contamination associated with the paraffin block or the reagents used during embedding and extraction. This issue is particularly relevant in low-biomass samples, such as solid-organ tissues, where exogenous DNA can be relatively more detectable ^[85]^. However, several studies have demonstrated that when appropriate methodological precautions are taken, contamination has a limited impact on biological conclusions. For instance, 16S rRNA analyses comparing FFPE tissue blocks, the surrounding paraffin, and extraction blanks revealed that the main source of bacterial contaminants originates from DNA extraction reagents rather than from the paraffin or the tissue itself ^[86]^. Similarly, other reports found negative 16S amplification in paraffin-only controls, supporting the notion that the contribution of paraffin to microbial signals is negligible ^[87]^. In our case, all liver biopsies were processed using a newly purchased commercial kit specifically designed for FFPE tissue microbiome extraction, and all samples originated from the same pathology center and were processed simultaneously under identical laboratory conditions. Under these standardized conditions, any background contamination would be expected to be evenly distributed across samples, minimizing its influence on intergroup comparisons. Therefore, although the possibility of trace contamination cannot be completely excluded, it is unlikely to have affected the differential patterns observed in this study.

In addition, several layers of stringency were incorporated into the bioinformatic workflow to further minimize potential artifacts and enhance the reliability of the results. Taxonomic and functional assignments were conducted using the DIAMOND–MEGAN6 pipeline ^[88]^ with conservative lowest common ancestor (LCA) parameters, including a minimal bit score of 87, a top-percent threshold of 5, and a minimum of three reads required for feature assignment. Moreover, reads mapped to *Deuterostomia*, *Panarthropoda*, or *Viridiplantae* were systematically excluded from downstream analyses to prevent spurious classifications arising from host or environmental DNA. These filtering steps substantially reduced the risk of false positives and improved the confidence in microbial profiling at both taxonomic and functional levels.

Likewise, statistically rigorous criteria were applied during differential abundance and functional enrichment analyses. The MaAsLin2 framework ^[20]^ was employed with CSS normalization to account for variable sequencing depth, and a negative binomial model was selected to properly handle the overdispersion typical of metagenomic count data. Only features detected in at least one-third of all samples and showing an adjusted q-value below 0.01 were retained as significant. Together, these restrictions ensured that the differences reported between groups reflected robust biological signals rather than stochastic variability or methodological noise, thereby strengthening the reproducibility and interpretability of our findings.

Finally, regarding the somatic variant-calling analysis, the absence of a matched “Panel of Normals” (PoN) for each sample represented another methodological limitation. Without a PoN, distinguishing true somatic mutations from germline variants or sequencing artifacts becomes more challenging, particularly in samples with low tumor purity such as FFPE liver tissue. To address this issue, a dual filtering strategy was implemented to maximize the specificity of variant detection. Variant calling was performed in “tumor-only” mode using *Mutect2* with a germline reference database to further exclude common population variants ^[29]^. Subsequently, variants were filtered using the *FilterMutectCalls* module from the GATK pipeline ^[28]^, followed by an additional stringent filtering step using *bcftools* ^[27]^ with conservative thresholds. Although the lack of a matched normal reference imposes inherent constraints on somatic variant interpretation, the combination of these conservative filtering parameters and the use of high-confidence germline databases effectively reduced false-positive calls and improved the reliability of the mutational profiles obtained in this study.

## CONCLUSION

In conclusion, this study demonstrates the feasibility and value of characterizing the hepatic microbiome as a true multi-kingdom ecosystem, integrating bacterial, fungal, protozoan, archaeal, helminth, and viral signatures from FFPE-derived shotgun metagenomes. By establishing a rigorously controlled and reproducible bioinformatic workflow tailored to low-biomass hepatic tissue —including conservative taxonomic assignment, multi-layer contamination control, kingdom-wide differential abundance testing, functional enrichment, and network inference— we provide a methodological framework that maximizes biological information while maintaining analytical stringency. Our findings show that functional alterations within the hepatic microbiota are more pronounced than global taxonomic shifts, and that specific microorganisms from every microbial kingdom differ across histological and genetic strata, underscoring the relevance of multi-kingdom analyses for understanding MASLD. These contributions highlight both the biological complexity of the hepatic niche and the importance of comprehensive, function-oriented metagenomic approaches for future biomarker discovery and mechanistic studies along the gut-liver axis.

## ACKNOWLEDGEMENTS

The authors would like to thank the patients for their cooperation.

## DATA AVAILABILITY STATEMENT

The dataset that support the findings of this study has been deposited in NCBI database as the sequence read archive (SRA) format (https://www.ncbi.nlm.nih.gov/sra/PRJNA1457056) under the accession number PRJNA1457056.

## Author contributions

Trinks J, Bustamante JP, and Penas-Steinhardt A designed and coordinated the study; Gadano A acquired funding; Marciano S, Casciato P, Narvaez A, Haddad L, and Gadano A recruited patients; Mascardi MF, Taussig R, Suarez B, and Bustamante JP processed the samples and performed sequencing; Mascardi MF, Privitera Signoretta I, and Trinks J analyzed data and interpreted the data; Mascardi MF and Trinks J wrote the manuscript; all authors approved the final version of the article.

**Supported by** PUE-CONICET grant N° 22920200100009CO, funds from IMTIB (CONICET-HIBA-IUHI), Research Council of Hospital Italiano of Buenos Aires grant, PIP-CONICET 2021–2023 grant 11220200100875CO, PICT-2020-Serie A-00788 and “Florencio Fiorini Foundation” grants to Dr Julieta Trinks; funds from Austral University for processing and sequencing of samples for microbiota analyses

## FOOTNOTES

### Institutional review board statement

This study was reviewed and approved by the Ethics Committee of Hospital Italiano de Buenos Aires.

### Informed consent statement

All participants have read and signed the informed consent of participation to this study.

### STROBE statement

The authors have read the STROBE Statement—checklist ofitems—and the manuscript was prepared and revised according to the STROBEStatement—checklist of items.

## SUPPLEMENTARY MATERIAL

**Supplementary Table 1. Sample-level taxonomic composition of hepatic metagenomes at the family, genus, and species levels**. Detailed taxonomic composition of liver tissue metagenomes at the family, genus, and species levels. The table reports the number of sequencing reads assigned to each taxon on a per-sample basis.

**Supplementary Table 2. Differentially abundant hepatic microbial taxa according to histological severity (NAS score).** Comprehensive list of hepatic microbial taxa showing statistically significant differences in abundance between low NAS and high NAS patient groups. Differential abundance analysis was performed at multiple taxonomic levels, and taxa are reported along with their corresponding fold-change coefficients and statistical significance metrics.

**Supplementary Table 3. Differentially abundant hepatic microbial taxa according to *PNPLA3* genotype.** Complete list of hepatic microbial taxa exhibiting statistically significant differential abundance between patients carrying the *PNPLA3* CC/CG genotype and those carrying the GG genotype. Results are reported across multiple taxonomic ranks, including fold-change estimates and associated statistical significance values.

**Supplementary Table 4. Somatic mutational burden metrics in FFPE liver biopsies of MASLD patients**. Per-sample summary of somatic mutational burden analysis performed on FFPE liver biopsy samples. The table reports the total number of high-confidence somatic variants identified per sample, sequencing coverage metrics, and normalized mutational burden estimates. The number of variants was normalized to the number of high-quality, uniquely aligned human reads (MAPQ ≥20), and expressed as variants per million reads to allow comparison across samples. Samples are annotated according to histological severity (low NAS vs. high NAS) and *PNPLA3* genotype (GG vs. CC/CG), and constitute the full dataset underlying the somatic mutational burden comparisons described in the Results section.

